# Cross-species developmental multiomics identifies *RHOJ* as a regulator of human-enriched cortical progenitor states

**DOI:** 10.64898/2026.11.21.768345

**Authors:** Yiming Sun, Kaimin Chen, Xiaoxia Yang, Benhui You, Fengyun Zhang, Xiaoyu Meng, Ming Li, Qing Cheng, Xiaoxuan Qi, Yuting Liu, Xiang Li, Yu Li, Tong Yan, Xuejiang Guo, Xin Luo, Cheng Li, Bing Su

**Author notes:** Corresponding author (B.S.); (C. L.); (X.L.). These authors contributed equally (Yiming Sun, Kaimin Chen).

## Abstract

Expansion of the primate cerebral cortex is associated with increased proliferation and diversification of neural progenitors, particularly outer radial glia (oRG), but the evolutionary developmental changes underlying these features remain unclear. Here, we integrate time-resolved, paired single-nucleus transcriptomic and chromatin-accessibility profiles from human, macaque and mouse fetal cortex to compare their radial glial developmental programs. Developmental alignment within a conserved corticogenic framework identified two human-enriched programs associated with oRG features and progenitor maintenance. We identified *RHOJ*, which encodes a conserved small GTPase, as an oRG-enriched gene with elevated expression in human cortex. Ectopic *RHOJ* expression in mouse fetal cortex increased the proportions of neural progenitors and cells with oRG-like morphology, and altered the relative abundance of *SATB2^+^* and *TBR1^+^* neuronal populations. An intronic *RHOJ* enhancer showed greater reporter activity for the human sequence than for orthologous non-human primate sequences, partly attributable to a human-specific substitution within a predicted AP-1 binding motif. In human cortical organoids, deletion of this enhancer reduced *RHOJ* expression, impaired organoid growth and altered progenitor and neuronal populations. These findings link human-enriched developmental programs to evolutionary modification of progenitor properties, supporting altered deployment of conserved genes as a potential contributor to cortical progenitor diversification.

## Introduction

Expansion of the cerebral cortex is a hallmark of human evolution^1–4^. Increased proliferative capacity of radial glia (RG) and expansion of the outer subventricular zone (oSVZ), which harbors abundant outer radial glia (oRG), are defining features of primate corticogenesis and have been implicated in the pronounced expansion of the human cortex^5–9^.

Besides of lineage-specific genes that have been shown to promote cortical progenitor proliferation and oRG expansion^10–16^, evolutionary changes in the regulation of genes shared across species provide another route to modifying progenitor development^17^. Understanding these changes requires a developmental perspective: RG transcriptional states evolve throughout corticogenesis, coupling progenitor properties to the sequential production of neuronal and glial populations^18^. Species differences in progenitor biology may therefore arise from changes in the timing, magnitude or coordination of conserved gene expression programs. However, it remains unclear which RG developmental programs diverge across species and how these differences contribute to the expansion and diversification of neural progenitor populations.

Here we generated paired single-nucleus transcriptomic and chromatin accessibility profiles from macaque and mouse cortex across multiple developmental stages, and integrated them with published human multiome data^19^. By aligning RG developmental stages across species, we distinguished shared temporal programs from human-enriched expression patterns associated with oRG features and progenitor maintenance. We then combined cross-species regulatory analyses with functional experiments in mouse cortex and human cortical organoids to investigate *RHOJ*, a conserved small GTPase gene enriched in human oRG. This approach connects developmental program differences to progenitor properties, providing a framework for investigating how altered regulation of conserved genes contributes to cortical progenitor diversification.

## Results

### Conserved temporal organization of cortical progenitor differentiation across species

To compare developmental features across species, we assembled a time-resolved atlas of paired single-nucleus transcriptome and chromatin accessibility profiles from developing human, macaque and mouse cortex (**Fig. 1a**). Newly generated 10x Genomics snMultiome data comprised 105,481 high-quality nuclei from macaque prefrontal cortex (PFC) at embryonic day (E) 50, E60, E70, E80, E90, E110 and postnatal day (P) 0, with two biological replicates per stage, and 69,160 nuclei from mouse rostral telencephalon or PFC at E11.5, E13.5, E14.5, E15.5, E16.5, E18.5 and P0, with two to three biological replicates per stage. We integrated these data with 58,816 nuclei from published human cortical multiome datasets spanning post-conception weeks (pcw) 11-35^19^ (**Methods; Extended Data Figs. 1-3**). This dataset captures key milestones of corticogenesis^20,21^.

**Fig. 1.**
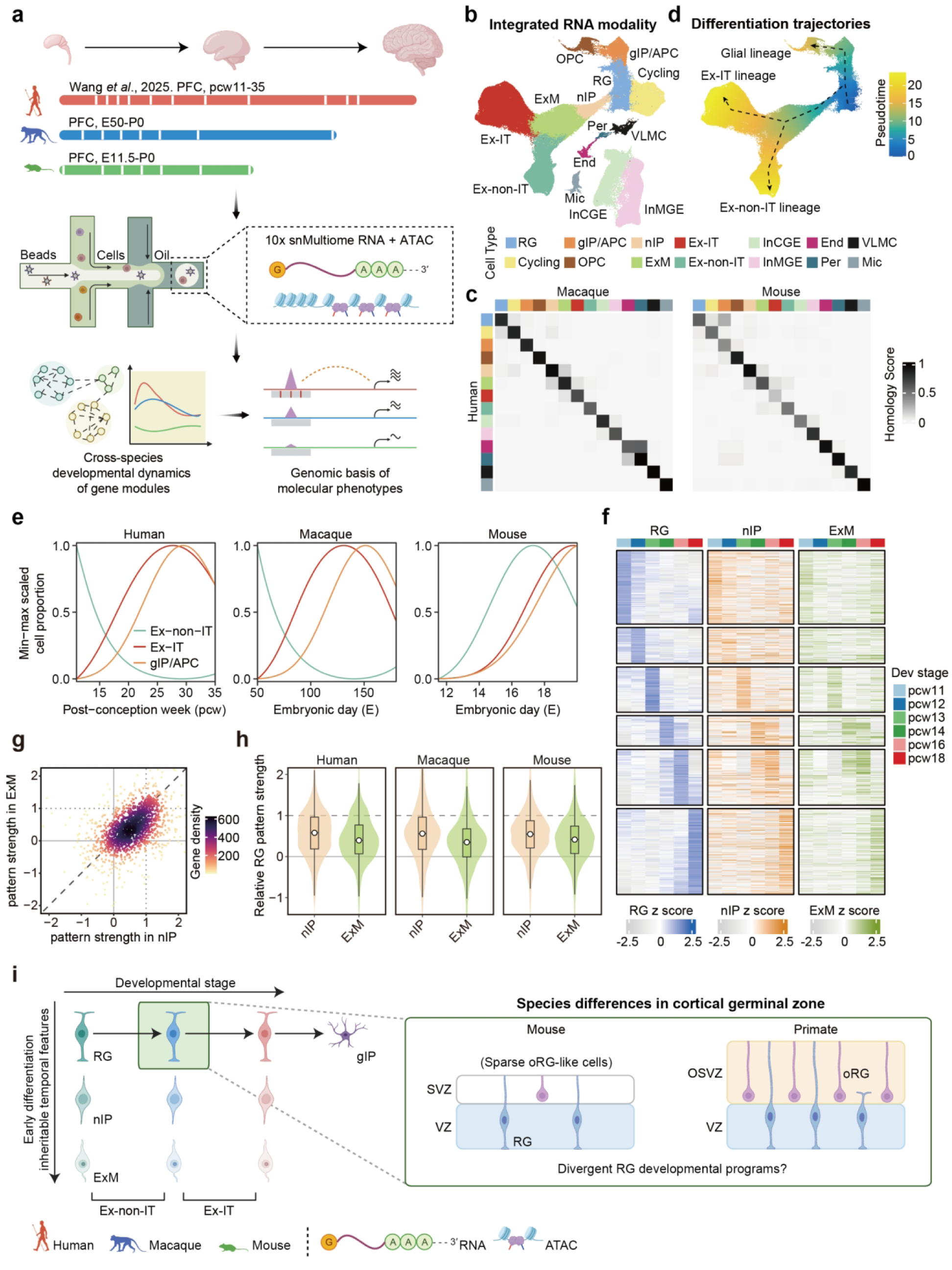
Conserved cellular and temporal organization of corticogenesis across species. **a**, Study design. Paired single-nucleus transcriptomic and chromatin accessibility profiles from developing macaque and mouse cortex were integrated with published human multiome data. Samples span post-conception weeks (pcw) 11–35 in human, embryonic day (E) 50 to postnatal day (P) 0 in macaque, and E11.5-P0 in mice. Cross-species analyses connect developmental gene expression programs to candidate regulatory differences. **b**, Uniform manifold approximation and projection (UMAP) of integrated transcriptomic profiles, colored by cell types. **c**, Correspondence of independently annotated human cell types with macaque (left) and mouse (right) cell types, quantified by co-clustering-based homology scores in the integrated dataset. **d**, Inferred developmental trajectories in the integrated transcriptomic embedding, colored by pseudotime. Dashed lines indicate trajectories connecting RG to excitatory neuronal and glial populations. **e**, Fitted developmental profiles of Ex-non-IT, Ex-IT and gIP/APC proportions, independently min-max scaled. Unscaled sample-level proportions, fitted curves and confidence intervals are shown in Extended Data Figs. 5-7. **f**, Temporal expression profiles of 1,741 temporally dynamic human RG genes in RG, nIP and ExM (genes with zero variance across stages in nIP or ExM were excluded). Genes are grouped and ordered by their stage of maximum expression in RG, and this row order and the RG-referenced numerical scale are retained across cell types. **g**, Relative strength of RG temporal expression patterns in human nIP and ExM. Each point represents a gene, colored by local gene density. Pattern strength is the regression slope of the downstream expression profile against the corresponding RG profile across matched developmental stages (Methods). Values between 0 and 1 indicate a weaker same-direction component than in RG; negative values indicate an opposite-direction component. The dashed diagonal indicates equal strength in nIP and ExM; dotted lines indicate the RG reference strength of 1. **h**, Distributions of relative RG temporal pattern strength in nIP and ExM, calculated separately for each species (human, 1,741 genes; macaque, 1,979 genes; mouse, 2,650 genes). **i**, Conceptual summary of temporal progenitor features and their partial persistence during differentiation, based on the framework of Telley et al. (2019)^18^, alongside differences in cortical germinal-zone organization. RG, radial glia; oRG, outer radial glia; gIP/APC, glial intermediate progenitors/astrocyte precursor cells; Cycling, cycling cells; nIP, neuronal intermediate progenitors; ExM, migrating excitatory neurons; Ex-IT, intratelencephalic-projecting excitatory neurons; Ex-non-IT, non-intratelencephalic-projecting excitatory neurons; InMGE and InCGE, medial and caudal ganglionic eminence-derived inhibitory neurons, respectively; OPC, oligodendrocyte precursor cells; End, endothelial cells; Per, pericytes; VLMC, vascular leptomeningeal cells; Mic, microglia; VZ, ventricular zone; SVZ, subventricular zone; oSVZ, outer subventricular zone.

Independent annotation identified corresponding broad cortical cell classes across the three species, supported by concordant marker expression and chromatin accessibility^22,23^ (**Fig. 1b, c**; **Extended Data Figs. 1f, g, 2f, g, 3f,g, and 4**). Trajectory inference recovered similar paths connecting radial glia (RG) to excitatory neuronal and glial populations (**Fig. 1d**; **Extended Data Figs. 5a, 6a and 7a**). Sampled cell-type proportions followed the expected progression from earlier enrichment of deep-layer-associated neuronal populations to later increases in other excitatory neuronal populations and glial progenitors^8,19,24–31^ (**Fig. 1e**; **Extended Data Figs. 5b, c, 6b,c, and 7b, c**). Differences in the shapes of these profiles were interpreted in the context of the distinct developmental windows sampled in each species.

We next asked whether temporal molecular features of RG were also detectable in downstream progenitor and neuronal states. Previous work in mice identified age-dependent progenitor transcriptional states that are partially retained during neuronal differentiation^18^. We identified temporally dynamic genes in RG of each species and compared their expression profiles with those in neuronal intermediate progenitors (nIP) and migrating excitatory neurons (ExM). Across all three species, nIP and ExM retained the same-direction component of RG temporal expression patterns, with pattern strength generally lower in nIP than in RG and further attenuated in ExM (**Fig. 1g, h**; **Extended Data Fig. 8**). Temporally dynamic accessibility peaks likewise showed partial persistence of RG-associated temporal features, although the extent differed across species and between modalities (**Extended Data Figs. 9-11**). These observations support shared temporal molecular organization across progenitor and differentiating neuronal states, without directly establishing lineage transmission.

Thus, broad cellular identities, developmental ordering and the partial persistence of RG temporal signatures were shared across species (**Fig. 1i**). This conserved organization provided a framework for investigating how RG developmental programs differ in species with markedly different progenitor populations and cortical architectures^9,32–34^.

### Developmental alignment reveals human-enriched radial glia programs

To resolve developmental programs within RG, we applied weighted gene co-expression network analysis (WGCNA)^35^ to temporally dynamic genes in human RG. Six modules, comprising 71-382 genes, captured early, transient and late expression programs (**Fig. 2a**; **Extended Data Fig. 12; Supplementary Table 1**). Early modules H_M1 and H_M2 were enriched for DNA replication and ribosome biogenesis (RiBi), respectively. Transient modules H_M3 and H_M4 were enriched for small GTPase signaling and stem cell population maintenance, whereas late modules H_M5 and H_M6 were enriched for cell communication and glial differentiation (**Fig. 2b, c**). Analysis of RG subtypes^17^ in an independent human dataset^36^ and ATAC-derived gene activity provided additional support for the temporal organization of these modules (**Extended Data Fig. 13**).

**Fig. 2.**
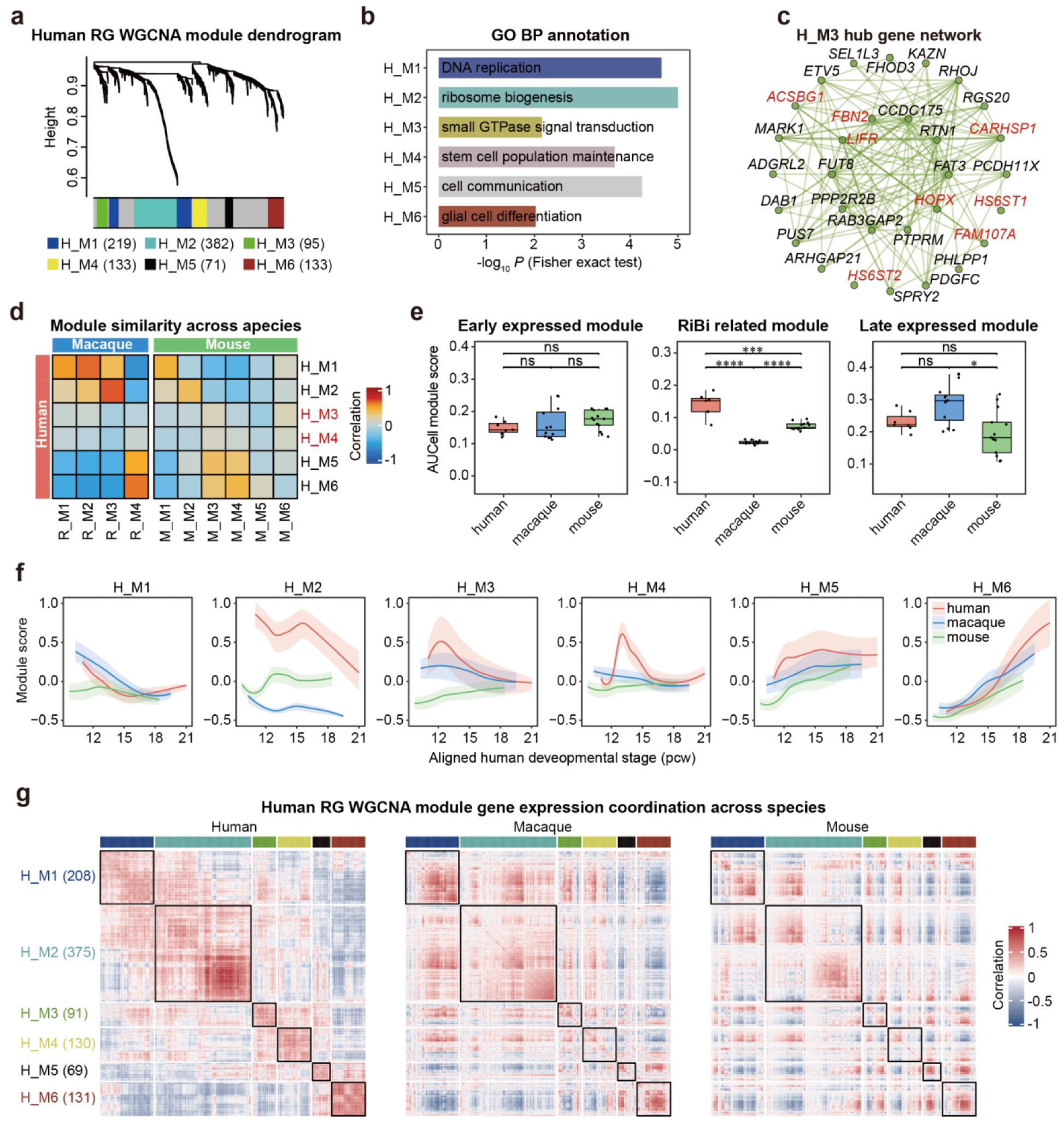
Conserved and human-enriched radial glia developmental programs. **a**, Hierarchical clustering of temporally dynamic human RG genes by weighted gene co-expression network analysis (WGCNA). The colored bar indicates module assignments; gray denotes unassigned genes. Numbers in parentheses indicate module sizes. **b,** Representative enriched Gene Ontology (GO) biological process terms for the six human modules. Bars show -log₁₀ P from Fisher’s exact tests. Additional terms are shown in Extended Data Fig. 12c. **c**, Co-expression network of 30 H_M3 hub genes. Previously reported oRG marker genes are labelled in red. **d**, Pearson correlations between module-membership (kME) profiles of human and macaque (left) or mouse (right) RG modules in a common gene space. H_M3 and H_M4 are labelled in red. H, human; R, rhesus macaque; M, mouse. **e,** AUCell scores for early, ribosome biogenesis (RiBi)-associated and late RG programs. Gene sets combine corresponding modules and are restricted to genes shared across species: early, H_M1/R_M1/R_M2/M_M1; RiBi, H_M2/R_M3/M_M2; late, H_M5/H_M6/R_M4/M_M3/M_M4. Each point represents the median cellular score within one source sample (human, n = 7; macaque, n = 12; mouse, n = 13). Statistical significance was assessed using two-sided Wilcoxon rank-sum tests with Benjamini-Hochberg correction. ns, adjusted P ≥ 0.05; *, adjusted P < 0.05; **, adjusted P < 0.01; ***, adjusted P < 0.001; ****, adjusted P < 0.0001. **f,** Expression dynamics of human RG modules in all three species along the aligned human developmental timeline (Extended Data Fig. 17a, b). Expression was standardized gene-wise across species before calculating cellular scores weighted by human kME. Lines show locally estimated scatterplot smoothing (LOESS) fits, and shaded bands show separately smoothed stage-wise 25th and 75th percentiles of cellular scores. Human, red; macaque, blue; mouse, green. **g**, Gene-gene Pearson correlation matrices for human module members across RG metacells (human, 280; macaque, 778; mouse, 1,078). The same 1,004 shared genes are grouped by human module and independently clustered within each module for each species. Numbers beside modules indicate retained genes. RG, radial glia; oRG, outer radial glia; pcw, post-conception week.

Applying the same framework to macaque and mouse RG identified four and six modules, respectively (**Extended Data Figs. 14 and 15; Supplementary Table 1**). Comparison of module-membership profiles recovered corresponding early proliferation-associated, RiBi-associated, and late cell communication-associated programs (**Fig. 2d**). Early and late program scores were broadly comparable across species (**Fig. 2e**). RiBi-associated scores differed among species, but their ordering was not consistently reproduced in independent datasets **Data Fig. 16**). We therefore used shared genes from the conserved early and late programs to align RG developmental stages by transcriptomic similarity (**Methods; Extended Data Fig. 17a, b**), allowing species differences to be examined on an RG-based developmental timeline.

This alignment helped distinguish differences in sampled developmental windows from divergent program expression. Mouse modules M_M5 and M_M6, enriched for pattern specification and epithelial development, were most prominent at early stages, and related temporal profiles were also observed in an earlier human dataset^36^ (**Extended Data Figs. 15c and 17c-e**). By contrast, H_M3 and H_M4 showed human-enriched developmental expression profiles (**Fig. 2f**). Their constituent genes also showed weaker within-module co-expression in macaque and mouse (**Fig. 2g**).

The fitted H_M3 profile peaked near pcw12.5 in human, with a weaker elevation at corresponding macaque stages and no comparable peak in mouse (**Fig. 2f**). In human, module H_M3 was preferentially expressed in outer radial glia (oRG), contained established oRG markers including *HOPX*, *LIFR*, *FBN2* and *FAM107A*, and was enriched for genes associated with small GTPase signal transduction, cell motility and cell migration^9,39,45^ (**Fig. 2b,2c**; **Extended Data Figs. 12c and 13b**). Temporally, its expression peak broadly coincided with the early establishment of outer subventricular zone (oSVZ) during human corticogenesis^46^. These features associated H_M3 with cellular properties relevant to oRG organization.

H_M4 showed a sharper fitted peak around pcw13, with lower expression at corresponding macaque and mouse stages (**Fig. 2f**). Its transient increase was evident in both ventricular RG (vRG) and oRG cells (**Extended Data Fig. 13b**). Enrichment for stem cell population maintenance, cell cycle and sterol biosynthesis, together with the inclusion of *HES1*, *HES5*, *SOX2* and *STAT3*, associated this module with progenitor maintenance and metabolic support (**Fig. 2b**; **Extended Data Fig. 12c, d**).

These results identified two human-enriched temporal programs within a conserved RG developmental framework. One is associated with oRG features and cell behavior, and the other with progenitor maintenance. We next examined their constituent genes to identify candidates linking these program-level differences to progenitor function.

### *RHOJ* is enriched in human outer radial glia

We compared gene expression and transcription start site (TSS) accessibility in RG across species during the aligned peak windows of H_M3 and H_M4 (**Supplementary Table 2**). In H_M4, 63 genes, approximately 47% of the module, were upregulated in human relative to macaque and mouse, whereas a further 29% showed a primate-upregulated pattern (**Extended Data Fig. 18**). Fifteen of the 63 human-upregulated genes, including *JUN*, *HES1* and *SOX9*, also showed human-elevated expression in independent sc/snRNA-seq datasets (**Extended Data Fig. 19a**). TSS accessibility was broadly comparable across species for many H_M4 genes, indicating that their expression differences were not uniformly accompanied by promoter accessibility differences. In H_M3, 26 genes, approximately 27% of the module, were upregulated in human relative to macaque and mouse, and 38 genes, approximately 40%, were upregulated in both primates relative to mouse (**Fig. 3a**; **Extended Data Fig. 20a**).

**Fig. 3.**
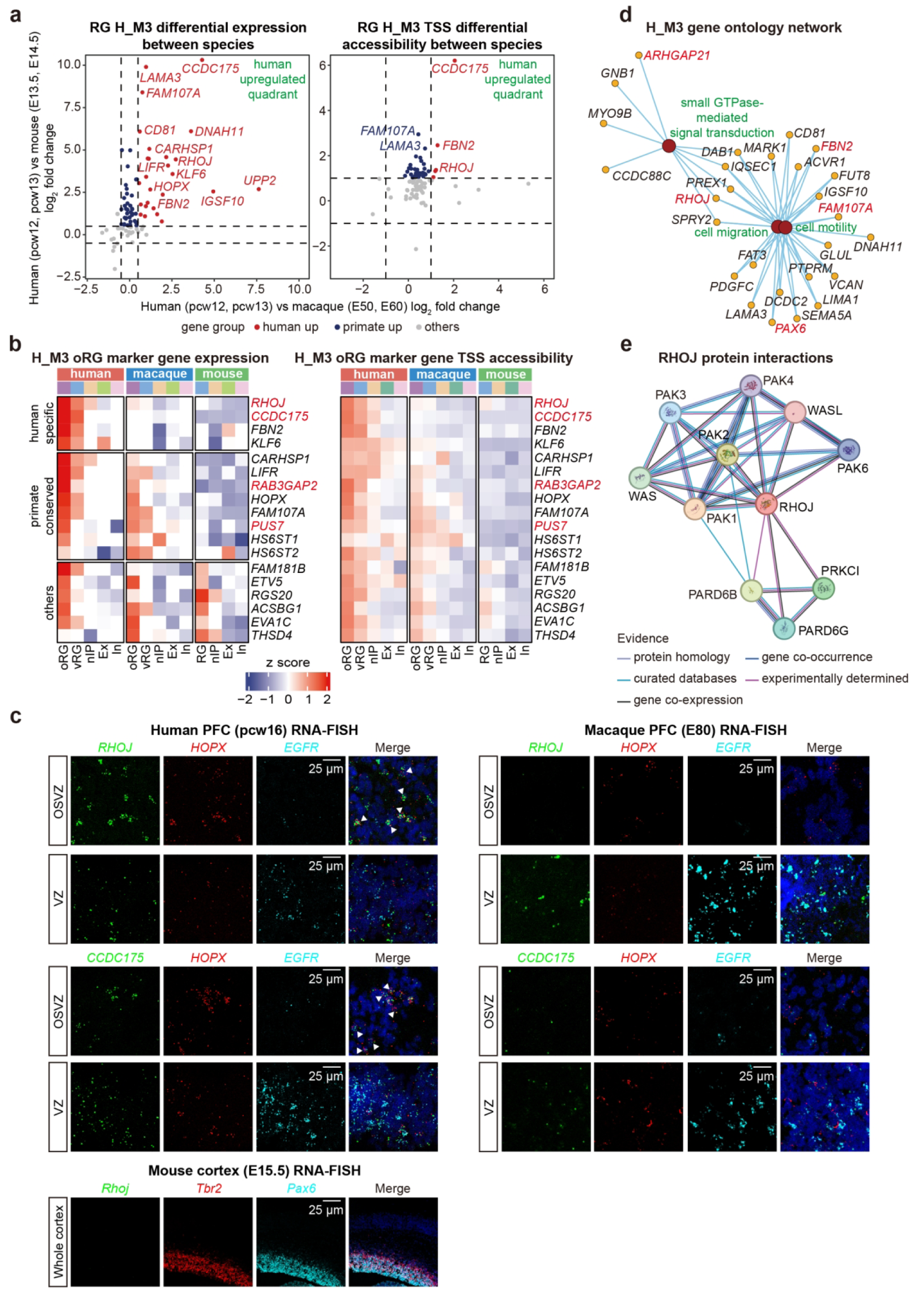
Identification of human-enriched outer radial glia genes. **a**, Cross-species differences in H_M3 gene expression (left) and transcription start site (TSS) accessibility (right). The x axis compares human pcw12-13 with macaque E50-60; the y axis compares human pcw12-13 with mouse E13.5-14.5. Each point represents a gene. Human-upregulated expression required log2 fold change > 0.5 and adjusted P < 0.01 in both comparisons, using Wilcoxon rank-sum tests. TSS accessibility was measured within ±2 kb of the TSS from a peak matrix normalized by Tn5 insertions around TSSs (ReadsInTSS); human-enriched accessibility required log2 fold change > 1 and adjusted P < 0.01 in both comparisons. Dashed lines indicate fold-change thresholds. Red, human-upregulated; blue, primate-upregulated; grey, other genes, classified separately for RNA and accessibility. **b**, Expression and TSS accessibility of H_M3 oRG-associated genes across species and cell types. Colors show gene-wise z scores across the displayed species and cell types, calculated separately for each modality. RHOJ, CCDC175, RAB3GAP2 and PUS7 are labelled in red. **c**, Representative RNAscope RNA-FISH images showing RHOJ or CCDC175 (green), HOPX (red) and EGFR (cyan) in human pcw16 and macaque E80 prefrontal cortex. Mouse E15.5 cortex shows Rhoj (green), Tbr2 (red) and Pax6 (cyan). Arrowheads indicate co-expression of RHOJ or CCDC175 with HOPX. DAPI, blue. Scale bars, 25 µm. **d**, GO network linking H_M3 genes to small GTPase-mediated signal transduction, cell migration and cell motility. Dark red nodes indicate GO terms; yellow nodes indicate genes; edges connect genes to annotated terms. Selected oRG-associated genes are labelled in red. **e**, STRING network of known and predicted functional associations. Edge colours indicate supporting evidence categories, as shown in the figure. RG, radial glia; oRG, outer radial glia; vRG, ventricular radial glia; nIP, neuronal intermediate progenitors; Ex, excitatory neurons; In, inhibitory neurons; oSVZ, outer subventricular zone; VZ, ventricular zone; pcw, post-conception week; E, embryonic day.

Examination across cell types identified *RHOJ* and *CCDC175* as oRG-enriched genes with higher expression in human, whereas *RAB3GAP2* and *PUS7* showed primate-upregulated expression (**Fig. 3b**; **Extended Data Fig. 20b**). *CARHSP1*, previously reported as a human oRG marker^39^, was also enriched in macaque oRG, whereas *FBN2* expression was predominantly associated with human oRG^45^. *RHOJ*, *CCDC175* and *FBN2* additionally exhibited higher TSS accessibility in human (**Fig. 3a, b**).

Independent fetal cortical datasets supported *RHOJ* and *CCDC175* expression in human RG, and spatial transcriptomic data supported their localization to the human oSVZ^47,48^ (**Extended Data Fig. 20c**). We performed RNAscope analysis and detected *RHOJ* and *CCDC175* signals in human oSVZ cells co-expressing *HOPX*, with limited *EGFR* co-expression (**Fig. 3c**). Sparse signals were observed in macaque germinal zones, whereas *Rhoj* signals were virtually absent in mouse fetal cortex. These observations supported the association of *RHOJ* and *CCDC175* with human oRG.

*CCDC175* encodes a coiled-coil domain-containing (CCDC) protein. Although other CCDC proteins have been implicated in cell proliferation and centrosome organization, the biological function of *CCDC175* remains poorly characterized^49–51^. We prioritized *RHOJ* for functional analysis because it combined human-enriched expression, oRG localization and a potential connection to cell behavior. *RHOJ* encodes a CDC42-related small GTPase and was associated with cell migration in the H_M3 GO network (**Fig. 3d**), suggesting that *RHOJ* may influence neural progenitor populations through cytoskeletal regulation^52,53^. STRING analysis also identified known or predicted functional associations with the PAK family (**Fig. 3e**), with its members implicated in neurodevelopmental disorders, including macrocephaly and intellectual disability^54–58^. These findings nominated *RHOJ* as a conserved gene whose increased expression might alter cortical progenitor properties.

### Ectopic *RHOJ* expression increases neural progenitors and oRG-like cells

To test whether increased *RHOJ* expression can influence cortical progenitor development, we ectopically expressed human *RHOJ* in the mouse cortex by in utero electroporation (IUE) at E13.5 and examined brains at E15.5 (**Fig. 4a**; **Supplementary Table 3**). *RHOJ* expression increased the proportions of PAX6^+^ RG and TBR2^+^ nIP cells among EGFP^+^ cells (**Fig. 4b, c**). Following a 2-hr BrdU pulse before tissue collection, the fractions of BrdU^+^ and Ki67^+^ cells within the EGFP^+^ population were also higher than in controls (**Fig. 4d-f**). These results showed enrichment of progenitor and cycling-cell states in the electroporated populations.

**Fig. 4.**
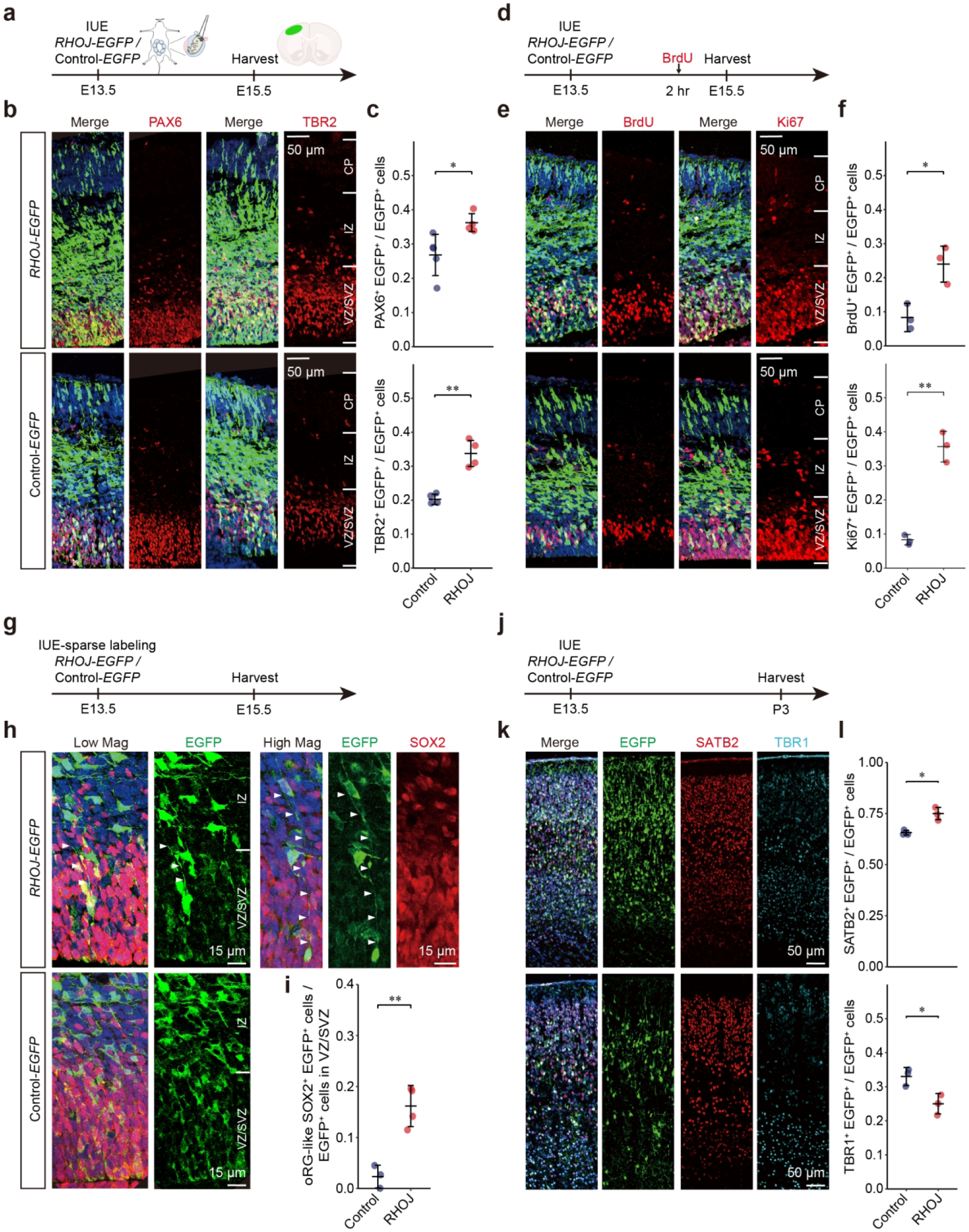
Ectopic *RHOJ* expression increases cycling-cell and oRG-like-cell fractions in mouse cortex. **a**, In utero electroporation (IUE) of human RHOJ or control plasmids at E13.5, followed by tissue collection at E15.5. **b**, Representative E15.5 cortical images showing EGFP (green) and PAX6 or TBR2 (red). **c**, Proportions of PAX6+ and TBR2+ cells among EGFP+ cells (n = 4 mice per group). Statistical significance was assessed using a two-tailed unpaired t-test. Data are shown as mean ± s.d. **d**, BrdU pulse labeling. BrdU was administered 2 hr before tissue collection at E15.5 following IUE at E13.5. **e**, Representative images showing EGFP (green) with BrdU or Ki67 (red). **f**, Proportions of BrdU+ and Ki67+ cells among EGFP+ cells (n = 3 mice per group). **g**, Sparse-labelling IUE design for morphological analysis at E15.5. **h**, Representative images showing EGFP (green) and SOX2 (red). Arrowheads identify SOX2+ EGFP+ cells with a long basal process and no apparent ventricular attachment, classified as oRG-like. **i**, Fraction of oRG-like SOX2+ EGFP+ cells among EGFP+ cells in the VZ/SVZ (n = 3 mice per group). **j**, IUE at E13.5 followed by tissue collection at P3 to examine neuronal output. **k**, Representative P3 images showing EGFP (green), SATB2 (red) and TBR1 (cyan). **l**, Proportions of SATB2+ and TBR1+ cells among EGFP+ cells (n = 3 mice per group). In c, f, i and l, points represent mice; data are mean ± s.d.; comparisons use two-sided unpaired t-tests. *, P < 0.05; **, P < 0.01. DAPI, blue. Scale bars, 50 µm (b, e and k) and 15 µm (h). CP, cortical plate; IZ, intermediate zone; VZ, ventricular zone; SVZ, subventricular zone; oRG, outer radial glia; E, embryonic day; P, postnatal day.

We next used lower-concentration plasmids for sparse labeling to assess individual progenitor morphology (**Fig. 4g**). A subset of SOX2^+^ EGFP^+^ cells displayed a long basal process directed toward the pial surface and no apparent apical attachment to the ventricle, resembling oRG morphology^9^ (**Fig. 4h**). The proportion of these oRG-like cells among EGFP^+^ cells in the VZ/SVZ was higher after *RHOJ* expression than in controls (**Fig. 4i**). Thus, increased *RHOJ* expression affected both the representation of cycling progenitors and the prevalence of progenitors with oRG-like morphology.

To assess later neuronal composition, we examined electroporated cortices at P3 (**Fig. 4j, k**). *RHOJ* expression increased the proportion of SATB2^+^ neurons and decreased the proportion of TBR1^+^ neurons among EGFP^+^ cells (**Fig. 4k, l**), suggesting a shift toward upper-layer neuronal output. This result mirrors the lineage bias of primate oRG cells, which are known to preferentially generate upper-layer neurons^7^.

Together, these results indicate that ectopic *RHOJ* expression can alter cortical progenitor states, morphology and subsequent neuronal output in mouse. They provided functional support for investigating increased *RHOJ* expression as a contributor to oRG generation and expansion during human corticogenesis.

### A human-specific substitution boosts *RHOJ* enhancer activity

We next investigated the regulatory basis of elevated *RHOJ* expression in human RG. Expression differences in many H_M3 and H_M4 genes were not accompanied by similarly pronounced TSS accessibility differences, motivating an examination of non-promoter regulatory regions. Human-enriched expression of transcription factors (TFs) such as *JUN* also suggested a possible contribution from the *trans*-regulatory environment (**Fig. 5a**).

**Fig. 5.**
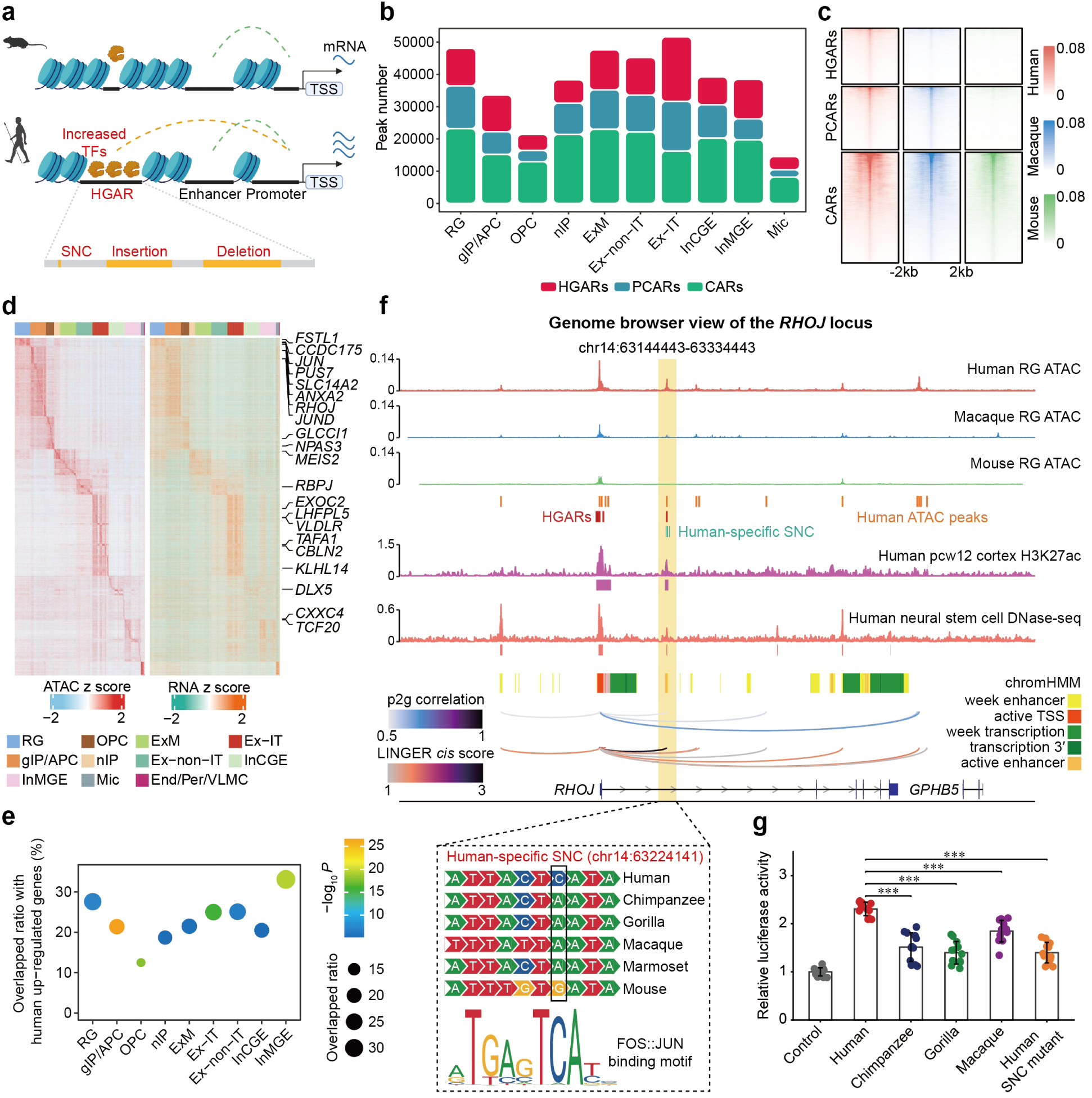
A human-specific substitution contributes to increased RHOJ enhancer reporter activity. **a**, Proposed contributions of cis-regulatory sequence changes and altered transcription factor (TF) availability to species differences in gene expression. **b**, Numbers of human-gained accessible regions (HGARs), primate-conserved accessible regions (PCARs) and conserved accessible regions (CARs) across cortical cell types. HGARs denote regions with higher accessibility in human than in macaque and mouse. **c**, ReadsInTSS-normalized Tn5 insertion signals within ±2 kb of region centers in human, macaque and mouse RG. Each row represents one region; row order is identical across species. Regions are grouped as HGARs, PCARs or CARs. **d**, Accessibility (left) and linked gene expression (right) for peak-gene pairs involving HGARs that overlap human evolution-associated regions or single-nucleotide changes (SNCs). Rows represent peak-gene pairs; columns represent human metacells, annotated by cell type above. Colors indicate row-wise z-scores. The displayed links were retained at correlation ≥ 0.20. **e**, Enrichment of human-upregulated genes among putative targets of HGARs overlapping human evolution-associated sequence changes. Dot size and position indicate the percentage of targets upregulated in human relative to both macaque and mouse; color indicates −log₁₀ P from one-sided Fisher’s exact tests. **f**, Genome browser view of the RHOJ locus, showing RG accessibility across species, HGARs, human-specific SNCs, human ATAC-seq peaks, published H3K27ac and DNase-seq signals, chromHMM states, peak-to-gene correlations and LINGER-predicted cis-regulatory links. The highlighted intronic region contains five human-specific SNCs. The alignment below shows the substitution at chr14:63224141 and a predicted FOS::JUN recognition motif. Coordinates refer to GRCh38. **g**, Luciferase reporter activities in SK-N-SH cells for the human candidate enhancer (chr14:63223858-63225232), its chimpanzee, gorilla and macaque orthologues, and the human sequence with the ancestral allele restored at chr14:63224141. Firefly activity was normalized to the co-transfected Renilla control (n = 4 per group). Bars show mean ± s.e.m. Comparisons use two-sided unpaired t-tests. *, P < 0.05; **, P < 0.01; ***, P < 0.001. RG, radial glia; TSS, transcription start site; cell-type abbreviations are defined in Fig. 1.

Using reciprocal LiftOver^59^, we constructed a set of accessible regions with one-to-one genomic correspondence across human, macaque and mouse (**Extended Data Fig. 21a, b**). Within this comparable sequence set, RG contained 11,697 human-gained accessible regions (HGARs), 13,235 primate-conserved accessible regions (PCARs) and 23,181 conserved accessible regions (CARs) (**Fig. 5b, c**; **Supplementary Table 4**). Here, HGARs denote regions with higher accessibility in human than in macaque and mouse. HGARs and PCARs were more frequently intronic and distal intergenic than CARs, consistent with reportedly greater divergence in non-promoter accessibility^60,61^ (**Extended Data Fig. 21c**).

RG HGARs were enriched for AP-1 related motifs, including FOS::JUN and FOSL2::JUND, whereas NFI-family and CTCF motifs were prominent in PCARs and CARs, respectively (**Extended Data Fig. 21d**). Within the cross-species-comparable sequence set, HGARs were enriched for human ancestor quickly evolved regions (HAQERs), but not significantly enriched for human accelerated regions (HARs) and human-specific structural variants, whereas regions of human-specific conserved deletions (hCONDELs) were underrepresented^62–65^ (**Extended Data Fig. 21e-h**). This contrast between HARs and HAQERs is consistent with findings from a human-chimpanzee tetraploid system, in which HAQERs, but not HARs, were enriched among regions with *cis*-regulated chromatin accessibility^66^. These observations suggest that human-lineage sequence changes do not necessarily correspond to increased chromatin accessibility and that different classes of sequence evolution may have distinct regulatory consequences.

Peak-to-gene correlations connected candidate accessibility differences to transcriptional differences (**Fig. 5d**; **Supplementary Table 5**). Putative target genes of HGARs overlapping human evolution-associated sequence changes were significantly enriched for human-upregulated genes across multiple cortical cell types (**Fig. 5e**). This analysis also recovered reported regulatory associations involving *MEIS2*, *NPAS3*, and the evolutionarily modified *CBLN2* E2 enhancer^67–69^.

At the *RHOJ* locus, we identified an intronic HGAR containing five annotated human-specific single-nucleotide changes (SNCs) (**Fig. 5f**). This region showed increased accessibility in human RG, overlapped fetal cortical H3K27ac signal and was annotated as a putative active enhancer by chromHMM^70,71^. One SNC, at chr14:63224141 in GRCh38, was predicted to create or strengthen a FOS::JUN binding motif. Together with human-enriched *JUN* expression in H_M4, this suggested a candidate connection between sequence evolution and the RG regulatory environment.

We compared the human enhancer sequence (chr14:63223858-63225232) with its chimpanzee, gorilla and macaque orthologues using luciferase assays in SK-N-SH cells. The human sequence drove higher reporter activity than all three non-human primate sequences. Reverting the SNC at chr14:63224141 to the ancestral allele reduced the activity of the human construct (**Fig. 5g**). Thus, this human-specific substitution contributed to enhanced reporter activity of the human sequence, identifying a candidate *cis*-regulatory mechanism for increased *RHOJ* expression. The predicted AP-1 interaction remained a mechanistic hypothesis.

### Deletion within the *RHOJ* enhancer impairs progenitor development in human cortical organoids

To test whether the candidate enhancer has a function in its endogenous genomic context, we deleted a region within it in H9 human embryonic stem cells using CRISPR/Cas9 and differentiated two independently derived clones, Enh-KO #1 and Enh-KO #2, into cortical organoids alongside wild-type controls (**Fig. 6a**; **Supplementary Table 6**). Genomic PCR, junction sequencing and TA cloning confirmed biallelic deletion, with no detectable sequence changes at the examined predicted off-target sites (**Fig. 6b**; **Extended Data Fig. 22a-c**). At day 50 (D50), *RHOJ* expression was lower in enhancer-deletion organoids, as assessed by qPCR (**Fig. 6c**).

**Fig. 6.**
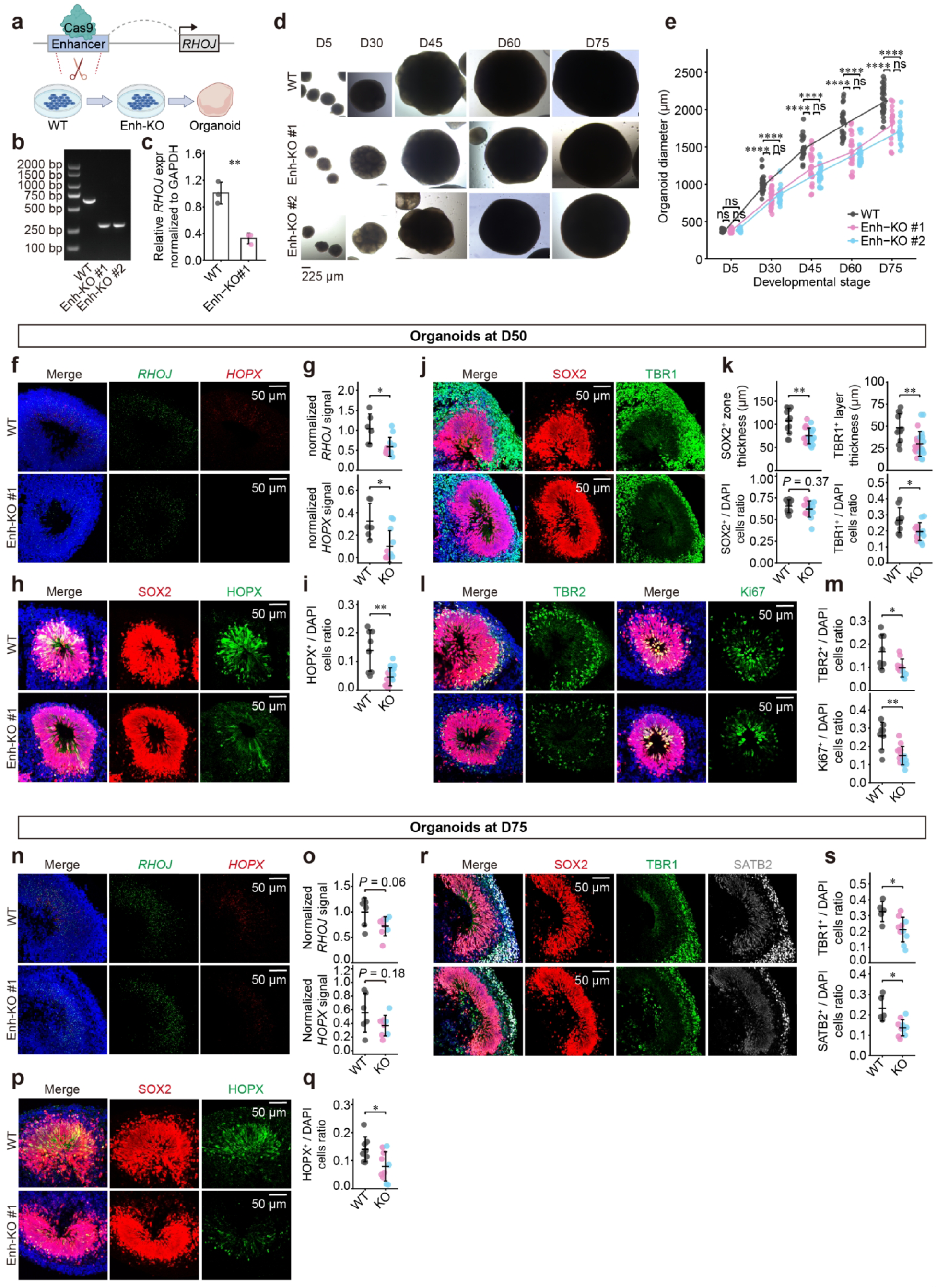
Deletion within the RHOJ enhancer alters progenitor development in human cortical organoids. **a**, CRISPR/Cas9-mediated deletion within the RHOJ enhancer in H9 human embryonic stem cells, followed by cortical organoid differentiation. **b**, Genomic PCR of wild-type (WT) cells and two independently derived deletion clones, Enh-KO #1 and Enh-KO #2. **c**, Relative RHOJ expression in D50 WT and Enh-KO #1 organoids, measured by qPCR and normalized to GAPDH (n = 3 per group). **d**, Representative bright-field images of organoids at D5, D30, D45, D60 and D75. Scale bar, 225 µm. **e**, Organoid diameters at the indicated stages. Each point represents one organoid. Numbers for WT, Enh-KO #1 and Enh-KO #2, respectively: D5, 33/21/31; D30, 40/28/24; D45, 20/25/27; D60, 26/23/28; D75, 27/17/17. Lines connect group means. **f**, Representative RNA-FISH images from Enh-KO #1 showing RHOJ (green) and HOPX (red). **g**, Normalized RNA-FISH signals, quantified as puncta per DAPI+ nucleus (three organoids and 5-6 neural rosette areas per cell line). **h**, Representative D50 immunofluorescence images from Enh-KO #1 showing SOX2 (red) and HOPX (green). **i**, HOPX+ fraction among DAPI+ cells (three organoids and 6-8 rosette areas per cell line). **j**, Representative D50 immunofluorescence images from Enh-KO #1 showing SOX2 (red) and TBR1 (green). **k**, SOX2+ progenitor zone and TBR1+ neuronal layer thickness, and SOX2+ and TBR1+ fractions among DAPI+ cells (three organoids and 6-8 rosette areas per cell line). **l**, Representative D50 images from Enh-KO #1 showing SOX2 (red) with TBR2 or Ki67 (green). **m**, TBR2+ and Ki67+ fractions among DAPI+ cells (four WT organoids and three organoids per deletion clone; 8-12 rosette areas per cell line). **n**, Representative D75 RNA-FISH images from Enh-KO #1 showing RHOJ (green) and HOPX (red). **o**, Normalized RNA-FISH signals, quantified as puncta per DAPI+ nucleus (three WT, three Enh-KO #1 and four Enh-KO #2 organoids; 4-8 rosette areas per cell line). **p**, Representative D75 images from Enh-KO #1 showing SOX2 (red) and HOPX (green). **q**, HOPX+ fraction among DAPI+ cells (three WT, three Enh-KO #1 and four Enh-KO #2 organoids; 4-8 rosette areas per cell line). **r**, Representative D75 images from Enh-KO #1 showing SOX2 (red), TBR1 (green) and SATB2 (gray). **s**, TBR1+ and SATB2+ fractions among DAPI+ cells (three WT, three Enh-KO #1 and four Enh-KO #2 organoids; 5 rosette areas per cell line). Representative cellular images show WT and Enh-KO #1; corresponding Enh-KO #2 images are in Extended Data Fig. 22. In g, i, k, m, o, q and s, each point represents one neural rosette area. Measurements from both deletion clones are grouped as Enh-KO. Grey, WT; pink, Enh-KO #1; cyan, Enh-KO #2. Data are mean ± s.d. Comparisons use two-sided unpaired t-tests. ns, not significant; *, P < 0.05; **, P < 0.01; ***, P < 0.001; ****, P < 0.0001. DAPI, blue. Scale bars, 50 µm (f, h, j, l, n, p and r). D, day of differentiation.

Although organoid diameters were comparable at D5, both deletion clones produced smaller organoids from D30 onward, with the difference persisting through D75 (**Fig. 6d, e**). At D50, Enh-KO#1organoids showed reduced *RHOJ* and *HOPX* RNA signals and a lower proportion of HOPX^+^ cells in neural rosette areas (**Fig. 6f-i**). The SOX2^+^ progenitor zone and TBR1^+^ neuronal layer were thinner, and the proportion of TBR1^+^ cells was lower, whereas the SOX2^+^ cell fraction was not significantly altered (**Fig. 6j, k**).

Fractions of TBR2^+^ intermediate progenitors and Ki67^+^ cycling cells were also reduced (**Fig. 6l, m**). Similar cellular phenotypes were observed in the independently derived Enh-KO #2 clone (**Extended Data Fig. 22d**).

At D75, enhancer-deletion organoids retained a lower proportion of HOPX^+^ cells and of both TBR1^+^ and SATB2^+^ neuronal populations (**Fig. 6p–s**; **Extended Data Fig. 22e**). *RHOJ* and *HOPX* RNA-FISH signals showed downward trends at this stage, although these differences were not statistically significant (**Fig. 6n, o**). The cellular effects of the deletion therefore persisted into later organoid development, despite weaker evidence for expression differences at D75.

These results supported an endogenous role for the tested enhancer region in *RHOJ* expression and cortical progenitor development. Combined with the reporter assays and mouse gain-of-function experiments, they connected a candidate human regulatory change to a gene capable of modifying progenitor properties. The contribution of the individual SNC in its endogenous context, and the extent to which *RHOJ* mediates the deletion phenotype, remain to be established. Together, the findings support regulatory modification of a conserved gene as a potential route to diversification of cortical progenitor development.

## Discussion

Our study links species differences in the temporal organization of radial glia development to the function and regulation of a conserved gene. Within a broadly shared framework of corticogenesis, developmental alignment identified two human-enriched RG developmental programs associated with oRG features and progenitor maintenance. These programs differed across species in both expression magnitude and within-module co-expression, suggesting that cortical progenitor diversification may involve changes in the coordination of conserved genes. Functional analysis of *RHOJ*, an oRG-enriched member of one program, connected increased expression to altered progenitor properties, while enhancer assays identified a human-specific substitution contributing to increased regulatory activity.

A developmental perspective is essential for interpreting such species differences. Comparisons at unmatched stages can conflate evolutionary divergence with differences in developmental timing or sampling coverage. By using shared early and late RG programs to align developmental progression, we distinguished human-enriched H_M3 and H_M4 profiles from early mouse programs (M_M5 and M_M6) whose apparent species specificity could partly reflect the developmental windows sampled. The weaker co-expression of H_M3 and H_M4 genes in macaque and mouse further suggests that differences extend beyond the abundance of individual transcripts to their temporal coordination. Nevertheless, these modules describe statistical relationships rather than established regulatory circuits, and their cross-species differences may also reflect variation in RG subtype composition. Comparisons within matched progenitor states will be important for separating changes in cell abundance from changes in gene regulation within cells.

The functional properties of H_M3 and H_M4 suggest two potentially complementary aspects of progenitor expansion. H_M3 was associated with oRG identity, small GTPase signaling and cell motility, whereas H_M4 was enriched for progenitor maintenance, cell cycle and sterol biosynthesis. Expansion of an oRG population requires both the organization of progenitors outside the ventricular zone and the maintenance of their capacity to proliferate. These processes need not be coupled: previous work showed that mTOR signaling affects human oRG morphology and migration through CDC42 without detectable effects on proliferation or cell fate^72^. The association of *KLF6*, an H_M3 member, with a cholesterol-related program in human oRG suggests a possible connection to the metabolic functions represented in H_M4^73^. Similarly, *JUN* and *JUND* in H_M4 and a predicted AP-1 motif in the *RHOJ* enhancer nominate a candidate regulatory link between the modules. These observations motivate a model in which changes in progenitor behavior, maintenance and metabolic support act together, although functional interactions between the programs remain to be tested.

Within H_M3, *RHOJ* provides an experimentally supported connection between human-enriched expression and progenitor properties. Ectopic *RHOJ* expression in mouse cortex increased progenitor proliferation and the proportion of cells with oRG-like morphology, accompanied by a shift toward SATB2^+^ neuronal output. These findings indicate that increased *RHOJ* expression can affect both progenitor states and cellular organization. In endothelial cells, *RHOJ* regulates focal adhesion disassembly through the GIT-PIX complex, which includes β-PIX, encoded by *ARHGEF7*^74^. An analogous effect on adhesion or cytoskeletal dynamics could influence progenitor attachment, basal processes or positioning. However, this mechanism has not been established in RG, and live imaging and lineage analysis will be needed to determine whether *RHOJ* alters delamination, mitotic behavior, self-renewal or neuronal production.

Other findings support the broader relevance of small GTPase-associated regulation to neural development. *RAB3GAP2*, another H_M3 member, showed primate-upregulated expression, and its disruption in human fetal brain-derived organoids has been reported to reduce progenitor populations while increasing the neuronal fraction^75^. Small GTPase-associated factors have also been implicated in evolutionary changes in neuronal maturation, including *ARHGEF7*, *RASGRF2* and the human-specific duplicate *SRGAP2C*^76,77^. These examples concern distinct developmental processes, but collectively suggest that modification of signaling systems controlling cell structure and behavior can contribute to neural diversification. The *RHOJ* results extend this possibility to cortical progenitors without implying a shared downstream mechanism.

The regulatory analyses provide a candidate evolutionary basis for increased *RHOJ* expression. The human intronic enhancer sequence showed greater reporter activity than its chimpanzee, gorilla and macaque orthologs, and reverting one human-specific substitution to the ancestral allele reduced this activity. Deletion within the enhancer lowered *RHOJ* expression and altered progenitor-associated and neuronal populations in human cortical organoids. These experiments establish distinct links: the substitution contributes to reporter activity, and the enhancer region has an endogenous developmental function. Together with the mouse gain-of-function results, they support a model in which altered regulation of *RHOJ* contributes to progenitor diversification. Establishing the complete causal chain will require endogenous ancestral-allele replacement and rescue of the enhancer-deletion phenotype by restoring *RHOJ* expression. AP-1 binding and perturbation experiments will likewise be necessary to determine whether the predicted motif mediates the sequence effect and interacts with the human-enriched *trans*-regulatory environment.

At the genome-wide level, the differing associations of HGARs with HAQERs, HARs and other evolutionary annotations indicate that accelerated sequence evolution and increased chromatin accessibility are not interchangeable signatures. These comparisons are restricted to regions retained by cross-species mapping and therefore do not capture the full contribution of lineage-specific sequence gains, deletions or structural changes. Likewise, similar TSS accessibility despite divergent expression does not by itself establish distal enhancer regulation, because TF activity, chromatin interactions and post-transcriptional processes may also contribute. The *RHOJ* experiments illustrate how comparative accessibility and expression analyses can prioritize a regulatory locus for functional testing.

Several limitations qualify the broader evolutionary interpretation. Developmental alignment reduces stage mismatch but cannot eliminate differences in sampled regions, donor characteristics or technical factors, and an RG-derived timeline need not align every cortical cell population equally well. The higher RiBi-associated expression observed in human was not consistently reproduced in independent datasets and should therefore remain a provisional observation, despite prior evidence linking mTOR, MYC and ribosome biogenesis to progenitor development^75^. In addition, static oRG-like morphology in mouse and HOPX-associated phenotypes in organoids do not establish the full functional identity or lineage potential of native human oRG. Neither system directly tests evolutionary changes in cortical size or architecture.

Our findings support a model in which cortical progenitor diversification can arise through altered deployment of conserved genes within stage-dependent developmental programs. *RHOJ* offers a functional example connecting human-enriched expression, progenitor properties and an evolutionarily modified regulatory element. This complements mechanisms involving lineage-specific genes and highlights temporal coordination and *cis*-regulatory modification as potential contributors to the development of an expanded cortical progenitor system.

## Methods

### Sample collection and ethical approval

All animal procedures were performed in accordance with international standards and were approved by the Institutional Animal Care and Use Committee of the Kunming Institute of Zoology, Chinese Academy of Sciences (approval number: SMKX-SQ2021-091). Macaque (*Macaca mulatta*) samples were obtained from the Primate Research Center of the Kunming Institute of Zoology, Chinese Academy of Sciences, a facility accredited by the Association for Assessment and Accreditation of Laboratory Animal Care (AAALAC) International. Brain specimens were collected from 14 macaque embryos via cesarean section at 7 developmental stages: embryonic day (E) 50, E60, E70, E80, E90, E110, and postnatal day (P) 0 (n = 2 biological replicates per stage). The prefrontal cortex (PFC) was dissected from each specimen and cryopreserved at −80°C. No neuropathological abnormalities were observed in the specimens. These samples were utilized for 10x single-nucleus multiome (10x snMultiome) library construction and RNA fluorescence in situ hybridization (RNA-FISH).

Pregnant C57BL/6J mice (*Mus musculus*) were obtained from the Animal Center of the Kunming Institute of Zoology, Chinese Academy of Sciences. A total of 15 embryonic brains were collected across 7 developmental stages (E11.5, E13.5, E14.5, E15.5, E16.5, E18.5, and P0), comprising 3 biological replicates for E15.5 and 2 replicates for all other stages. The rostral telencephalon (due to the limited size of the developing brain at E11.5) or PFC was dissected for 10x snMultiome library construction and RNA-FISH. Pregnant ICR mice, obtained from GemPharmatech Co., Ltd., were used for *in utero* electroporation (IUE) experiments.

Human (*Homo sapiens*) fetal brain tissue at pcw16 was collected for RNA-FISH analysis. Research protocols were approved by the Reproductive Research Ethics Committee of Nanjing Maternity and Child Health Care Hospital (approval number: 2023key-085). Informed consent was obtained from all donors for the use of samples in research, with no direct medical or financial benefit provided to the donors or other individuals.

### 10x snMultiome library construction and sequencing

Single-nucleus library preparation was performed using a unified protocol for both macaque and mouse samples. Frozen tissue samples were ground into a fine powder in liquid nitrogen using a mortar and pestle. All buffers were freshly prepared and maintained on ice. Isolated nuclei were resuspended in 500 μL of diluted Nuclei Buffer (1× Nuclei Buffer [PN-2000207, 10x Genomics], 1 mM DTT, 1 U/mL RNase inhibitor, and nuclease-free water) and incubated on ice. Following washing and centrifugation steps, nuclei were resuspended and diluted to the optimal concentration. Nuclei quantity and quality were assessed using trypan blue staining and a Luna automated fluorescence cell counter, with a target recovery of 10,000 nuclei per library. Libraries were constructed according to the Chromium Next GEM Single Cell Multiome ATAC + Gene Expression User Guide (CG000338, 10x Genomics) and sequenced on an Illumina NovaSeq 6000 platform following the manufacturer’s recommendations.

### 10x snMultiome sequencing data preprocessing

Raw sequencing data were processed using the Cell Ranger ARC pipeline (v2.1.0). Species-specific reference indices (Mmul_10 for macaque and GRCm39 for mouse) were constructed using the ’cellranger-arc mkref’ command. The ’cellranger-arc count’ command was executed with the ’--include-introns’ parameter to capture intronic reads for single-nucleus analysis and jointly quantify gene expression and chromatin accessibility. For downstream analysis, the resulting gene expression matrices were analyzed using Seurat (v5.2.0)^78^, and the ATAC-seq fragment data were processed using ArchR (v1.0.3)^79^.

### 10x snMultiome data quality control and processing

To ensure cross-species comparability, identical quality control (QC) standards and processing workflows were applied to all human, macaque, and mouse datasets. The snMultiome data were split into paired transcriptomic (RNA-seq) and chromatin accessibility (ATAC-seq) modalities for independent QC processing.

For the transcriptomic modality, low-quality nuclei, defined as those with < 200 detected genes or a mitochondrial read percentage > 5%, were filtered out. Potential doublets (> 10,000 detected genes or > 50,000 total UMI counts) were also removed. For the chromatin accessibility modality, nuclei with < 1,000 unique fragments or a transcription start site (TSS) enrichment score < 4 were excluded. Only nuclei satisfying QC criteria in both modalities were retained for downstream analyses. To further identify potential doublets, Scrublet (v1.1.0) was applied to the transcriptomic data for each sample independently using default parameters^80^. Clusters in which more than 50% of cells were predicted to be doublets were identified as doublet clusters and removed.

Dimensionality reduction and clustering were performed as follows. For the transcriptomic data, the expression matrix was normalized, and the top 2,000 highly variable genes were identified.

Dimensionality reduction was performed using the ’RunPCA’ function to compute 50 principal components (PCs), followed by Uniform Manifold Approximation and Projection (UMAP) embedding and Louvain clustering (resolution = 0.8) based on the top 30 PCs. Cell types were annotated based on the expression of canonical marker genes, and these annotations were directly assigned to the paired chromatin accessibility profiles.

For the chromatin accessibility data, the ’addIterativeLSI’ function was used for feature selection and latent semantic indexing (LSI) dimensionality reduction (3 iterations, 20,000 features, top 50 dimensions). Clustering was performed using the ’addClusters’ function, and visualization was generated using the ’addUMAP’ function based on the top 15 LSI dimensions. Pseudo-bulk Tn5 insertion profiles (coverage matrices) were computed for each cell type using the ’addGroupCoverages’ function. Subsequently, reproducible peaks were identified using the ’addReproduciblePeakSet’ function, which invokes MACS2^81^ (maxPeaks = 150,000; *P* cutOff = 0.05). Finally, a peak matrix was generated using the ’addPeakMatrix’ function for downstream analyses.

### Cross-species data integration and cell-type similarity evaluation

Transcriptomic datasets from human, macaque, and mouse were individually annotated based on clustering results and the expression of known marker genes. To evaluate global conservation across species, we integrated the transcriptomic data using the Reciprocal PCA (RPCA) workflow in Seurat. First, one-to-one orthologous genes shared among human, macaque, and mouse were retrieved using the biomaRt package (Ensembl release 115)^82^. The datasets were subset to these shared genes and merged. The top 2,000 highly variable genes were identified using the ’vst’ selection method. Based on these features, integration anchors were identified by projecting datasets into a shared PCA space, and the data were subsequently integrated. After integration, cross-species cell-type similarity was quantified by calculating the co-clustering proportion of corresponding cell populations, as described in previous studies^22,23^:

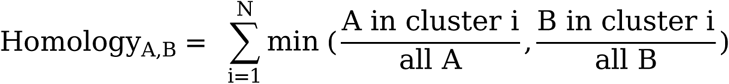

where A represents a specific cell type from species 1, B represents a cell type from species 2, N denotes the total number of clusters identified in the integrated dataset, and min represents the minimum function.

### Cortical cell differentiation trajectory analysis

To characterize cell state transitions and lineage progression across human, macaque, and mouse, we performed trajectory inference using Slingshot (v2.18.0) based on UMAP embeddings derived from the transcriptomic modality of the snMultiome data^83^. Inhibitory neurons and non-neural cell populations were excluded due to the incomplete capture of their differentiation lineages in the current dataset. Radial glia (RG) were designated as the root state, while late-stage excitatory neurons (Ex) and oligodendrocyte precursor cells (OPC) were defined as the terminal states.

### Developmental changes in cell-type composition

Cell-type proportions were calculated for each sample by dividing the number of nuclei assigned to a cell type by the total number of retained annotated nuclei. Developmental changes were fitted separately for each species and cell type using beta-binomial regression with a logit link in glmmTMB. The response comprised the numbers of nuclei belonging and not belonging to the cell type. Chronological age was centered and scaled within each species. P0 was assigned E180 in macaque and E20 in mouse. Predictions were restricted to the sampled age range. Confidence intervals were calculated as the fitted logit ±1.96 standard errors and transformed to proportions. For the temporal-output overview, each species-by-cell-type fitted curve was independently rescaled to a range of 0-1 using its predicted minimum and maximum.

### RG temporal expression patterns in downstream cell populations

We first excluded low-abundance transcripts (bottom 10% of average expression within RG cells) and subsequently retained only genes showing a significant association with sampling time points, as determined by the tradeSeq^84^ association test (*P* < 0.01). This yielded a final set of 1,784, 1,994, and 2,676 temporally dynamic genes for human, macaque, and mouse RG cells, respectively.

For each gene, the downstream cell type temporal profile was regressed on its RG profile across the observed stages. RG-referenced temporal pattern strength was calculated as:

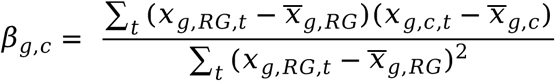

where x is log-transformed counts per million (CPM), t denotes stage and c denotes nIP or ExM. Heatmap z score were obtained by centering each gene separately within each cell type and dividing all three profiles by that gene’s RG temporal standard deviation. Genes were grouped by their stage of maximum RG expression and ordered within groups by their Spearman correlation with age. The same ordering and numerical scale were retained across cell types. Early-, intermediate-and late-maximum groups were summarized by their median scaled values at each stage.

### RG temporal chromatin accessibility patterns

Species-specific PeakMatrices were obtained from the ArchR objects. The feature universe was restricted to peaks overlapping reproducible RG peaks. Within each RG developmental stage, metacells were constructed using 30 neighboring nuclei in the first 30 iterative LSI dimensions. Counts and ReadsInTSS were summed over the constituent nuclei. Metacell peak counts were divided by their summed ReadsInTSS and multiplied by 10^6^.

To exclude low-signal peaks, those with the bottom 10% of average accessibility signal in RG cells were excluded. The normalized metacell peak signal matrix was transformed using log1p, and temporal differences were evaluated using limma with a stage-specific design matrix. All pairwise stage contrasts were fitted, followed by empirical Bayes moderation and Benjamini–Hochberg correction within each contrast. A peak was classified as temporally dynamic when at least one contrast had an absolute fitted difference > 1 and an adjusted *P* < 0.01.

For downstream comparisons, peak counts and ReadsInTSS were aggregated within each stage-by-cell-type group, normalized as above and log-transformed. RG-referenced strengths, heatmaps and grouped profiles were calculated using the same definitions as for RNA. Temporal peaks were selected independently in each species; peaks were not required to be orthologous. Comparisons of RNA and ATAC strength distributions were descriptive and did not treat genes and peaks as paired observations.

### RG weighted gene co-expression network analysis

Weighted gene co-expression network analysis (WGCNA) was performed on human, macaque, and mouse RG cells using hdWGCNA (v0.4.08) to identify developmental gene expression programs^35^. To prioritize biologically relevant features and reduce computational complexity, we refined the dataset to focus on temporally dynamic genes defined above.

To mitigate the sparsity inherent in single-nucleus data, metacells were constructed by aggregating mutually nearest neighbors in the RG PCA embeddings using the ’MetacellsByGroups’ function (k = 30; dims = 1:5). Co-expression networks were then built for each species using the ’ConstructNetwork’ function. Soft-power thresholds were selected to satisfy scale-free topology criteria following hdWGCNA guidelines, while other parameters were kept consistent to ensure cross-species comparability: human (soft power = 6; deepSplit = 3; mergeCutHeight = 0.4), macaque (soft power = 9; deepSplit = 3; mergeCutHeight = 0.4), and mouse (soft power = 8; deepSplit = 3; mergeCutHeight = 0.4).

### RG WGCNA module similarity across species

To quantify the conservation of gene co-expression networks across species, we evaluated the similarity of module membership (kME) patterns between corresponding modules. In WGCNA, the kME value represents the correlation between a gene’s expression profile and the module eigengene, serving as a robust measure of module membership. Given that homologous modules often differ in their gene composition across species, direct comparison requires a unified feature space. We therefore took the union of genes from the pair of modules being compared and recalculated the kME matrix for this combined gene set using the ’corSparse’ function in hdWGCNA. This step ensured that the module membership vectors were aligned to the same gene dimensions. Finally, module similarity was defined as the Pearson correlation coefficient between these recomputed kME vectors.

### RG developmental stage alignment across species

Given the substantial differences in the timing of cortical development among human, macaque, and mouse, aligning the developmental progression of RG cells across species was essential for a fair comparison of gene expression patterns.

This approach involved calculating the correlation of transcriptional profiles across developmental stages. We specifically selected WGCNA modules that were highly conserved across species and exhibited similar expression dynamics, hypothesizing that these modules reflect core developmental programs in RG cells. These selected modules included H_M1, H_M5, and H_M6 for human; R_M1, R_M2 and R_M4 for macaque; and M_M1, M_M3, and M_M4 for mouse. A unified feature set was established by taking the union of orthologous genes present in these modules. Using this gene set, we calculated pairwise Pearson correlation coefficients between the transcriptomic profiles of different developmental stages across species. Optimal alignment points were identified by pairing each developmental stage in one species with its most transcriptomically similar counterpart in the other based on maximal correlation. Using these matched data points, we fitted generalized additive models (GAMs; k = 3), weighted by the Pearson correlation coefficients, to model the continuous correspondence of developmental timing across species.

### WGCNA module dynamics across species and modalities

To characterize the developmental dynamics of WGCNA modules at the transcriptomic level across species, we first identified orthologous genes within each module shared among human, macaque, and mouse. Expression matrices for these genes were extracted from the Seurat object of each species, normalized, merged, and scaled feature-wise. For each cell, a composite module expression score was calculated as the weighted average of the scaled expression values, utilizing the module membership (kME) of each constituent gene as the weight. Module scores were plotted against native or aligned developmental age using local weighted regression (LOESS) with span 0.75 and degree 2. Shaded bands were obtained by calculating the 25th and 75th percentiles of cellular scores at each stage and smoothing each boundary separately.

To evaluate the consistency of module activity between transcriptomic and chromatin accessibility modalities, we extracted the gene activity score matrix (GeneScoreMatrix) from the RG ArchR object using the ’getMatrixFromProject’ function and the gene expression matrix from the RG Seurat object using the ’LayerData’ function. Both matrices were independently normalized and scaled feature-wise. For each modality, a composite module score was computed for each cell by weighting the scaled expression (or accessibility) values by the module membership (kME) of the constituent genes. Finally, the coordinated developmental dynamics of the modules between transcriptional and chromatin accessibility modalities were visualized using LOESS smoothing as described above.

### Cross-species chromatin accessibility comparison

To establish a unified feature set for cross-species comparison, we generated a set of consensus chromatin accessible regions. First, the complete set of reproducible peaks identified across all cell types was extracted from the ArchR project for each species. Using the UCSC LiftOver tool, chromatin accessible regions from macaque (Mmul_10) and mouse (GRCm39) were mapped onto the human reference genome (GRCh38)^59^. To ensure high sequence homology, we required the mismatch rate and sequence length difference between the original and mapped regions to be ≤ 10% for macaque and ≤ 40% for mouse. The ’bedtools merge’ command was then used to combine the human accessible regions with the mapped macaque and mouse regions^85^. To establish a one-to-one correspondence between species, these merged regions were reciprocally mapped back to the macaque and mouse genomes using the same filtering criteria, resulting in a final consensus peak set across the three species.

Next, this consensus peak set was added to the ArchR project for each species using the ’addPeakSet’ function. A new Tn5 insertion count matrix (PeakMatrix) corresponding to these genomic regions was computed using the ’addPeakMatrix’ function, with the ’ceiling’ parameter set to Inf. To mitigate data sparsity and balance the statistical power driven by large cell numbers, we employed a metacell strategy. Consistent with the hdWGCNA analysis, a k-nearest neighbor (KNN) network of RG cells was constructed based on the LSI embedding, and every 30 mutually nearest neighbors from the same sample were aggregated into a metacell. The merged chromatin accessibility matrix was normalized by the number of Tn5 insertions around transcription start sites (ReadsInTSS) to correct for sequencing depth and library quality. Signal intensities were further normalized by peak length to account for sequence length differences across species. Finally, differentially accessible peaks (DAPs) between species were identified using the two-sided Wilcoxon rank-sum test (|log_2_ fold change| > 1, *P* < 0.01).

### Transcriptional regulatory network analysis

For *cis*-regulatory network inference, we employed the built-in methods in ArchR and the LINGER framework (v1.0.0)^86^. In ArchR, peak-to-gene linkages were identified based on correlation analysis (correlation > 0.2, genomic window = ±250 kb from TSS) utilizing metacells derived from an LSI-based KNN graph. Default parameters were used for all other settings. For LINGER analysis, a KNN network was first constructed based on the PCA embedding of the transcriptomic data, and metacells were generated by aggregating every 100 mutually nearest neighbors from the same sample. The resulting merged gene expression and chromatin accessibility matrices were used to fine-tune a pre-trained neural network model. In this fine-tuned model, the Shapley value of each candidate chromatin accessible region quantified its contribution to the transcription of the target gene. Default parameters were used for LINGER.

### TF motif enrichment and binding site prediction

Sequences of RG consensus HGARs, PCARs and CARs were extracted from GRCh38. The 879 motifs in the JASPAR2024 CORE vertebrate non-redundant collection^87^ (v0.99.7) were scanned using ‘motifmatchr::matchMotifs’ at *P* < 5 × 10^-5^, using nucleotide frequencies estimated jointly from the retained sequences. Each interval was scored for the presence or absence of at least one match to each motif. Each accessibility class was compared with the other two classes combined. Intervals were stratified by GC-content deciles, length quintiles and promoter overlap. Within strata containing both comparison groups, motif enrichment was assessed using two-sided Cochran–Mantel–Haenszel tests without continuity correction. Mantel–Haenszel odds ratios summarized enrichment, and their log_2_ values were displayed. Benjamini–Hochberg correction was applied jointly across 879 motifs and 3 contrasts.

Representative motifs were selected from positive enrichments with adjusted *P* < 0.05, at least 20 motif-positive intervals in each comparison group and a frequency ≥ 2% in at least one group. Up to 6 motifs per class were selected by effect size, with overlapping selections combined.

To map precise TF binding sites and evaluate the impact of sequence evolution on TF binding affinity, we utilized the FIMO tool from the MEME Suite^88^. This analysis was performed using default parameters.

### Enrichment of human evolution-associated genomic regions

Human accelerated regions (HARs), human ancestor quickly evolved regions (HAQERs), hCONDEL-associated regions and human-specific structural variants (SVs) were obtained from published resources and represented in GRCh38 coordinates^62–65^. Overlap was defined as at least one shared base, irrespective of strand; multiple overlaps with the same annotation class counted once per accessible region.

Enrichment was assessed separately for RG and the nonredundant HGAR union across ten cell types. The RG universe comprised 80,086 consensus intervals tested in both human-to-macaque and human-to-mouse comparisons and overlapping human RG called peaks, including 11,697 HGARs. The all-cell-type universe comprised the union of eligible intervals across cell types (200,642 intervals), of which 65,915 were HGARs in at least one cell type. Within each scope, non-HGAR intervals formed the background. Full consensus-interval lengths were retained. Regions were stratified by chromosome, length quintile and promoter overlap, with length boundaries determined separately for each scope. Conditional permutations preserved the numbers of regions, HGARs and annotation-positive regions within every stratum.

Hypergeometric sampling generated 100,000 null HGAR-overlap counts. Two-sided *P* values were calculated as (1 + number of permutations with |N_null_ − E| ≥ |N_observed_ – E|)/100,001, where E is the conditional expected overlap count. *P* values were Benjamini–Hochberg-adjusted across the eight scope-annotation comparisons. Null distributions were displayed as overlap percentages; enrichment estimates were Mantel–Haenszel odds ratios with approximate 95% confidence intervals.

### *In utero* electroporation and BrdU labelling

IUE was performed on pregnant ICR mice at embryonic day 13.5 (E13.5). The coding sequence (CDS) of human *RHOJ* was cloned into the pCAGGS vector, with the empty pCAGGS vector serving as the negative control. To assess the impact of ectopic human *RHOJ* expression on RG cell proliferation and neurogenesis, plasmid DNA was prepared at a high concentration (1.5 µg/µL). In contrast, for morphological analysis requiring sparse labeling, the plasmid concentration was reduced to 0.2 µg/µL. In all cases, plasmid solutions were mixed with 0.1% Fast Green to visualize the injection.

Surgical procedures followed previously established protocols. Briefly, plasmid solutions were microinjected into the lateral ventricles of embryonic brains, and electroporation was delivered using a BTX830 electroporator with the following settings: five pulses of 35 V, 50 ms duration, with 950 ms intervals. Mice were maintained under isoflurane anesthesia throughout the procedure. After electroporation, embryos were returned to the abdominal cavity, and the incision was sutured. Mice were monitored until full recovery. Brain samples were collected at E15.5 and P3, with at least three biological replicates analyzed per group.

For cell proliferation analysis, 5-bromo-2′-deoxyuridine (BrdU; Sigma, B5002) was dissolved in PBS to prepare a 10 mg/mL stock solution. Pregnant mice were electroporated at E13.5 with plasmid DNA at 1.5 µg/µL. BrdU was injected intraperitoneally (50 mg/kg) at E15.5, 2-hr prior to sample collection, to label proliferating cells in the S phase.

### RNA fluorescence *in situ* hybridization (RNA-FISH)

RNA-FISH was performed on 10 μm tissue sections from human, macaque, and mouse brain samples, and 20 μm sections from human cortical organoids using the RNAscope^®^ Multiplex Fluorescent Reagent Kit v2 (Advanced Cell Diagnostics, ACD). Hybridization and signal detection were conducted according to the manufacturer’s standard instructions. For human and macaque samples, identical probe sets were employed to examine the co-expression of *RHOJ* and *CCDC175* with the oRG marker *HOPX* and the gIP marker *EGFR*. In mouse samples, the co-localization of *Rhoj* with the neural progenitor markers *Pax6* and *Eomes* was assessed using mouse-specific probes.

### Immunofluorescence staining

Brain tissues were fixed in 4% paraformaldehyde (PFA) for 12-hr and subsequently cryoprotected by sequential immersion in 15% and 30% sucrose solutions at 4°C. Mouse brain sections of 10 μm thickness and cortical organoid sections of 20 μm thickness were prepared for further use.

For immunofluorescence staining, sections were subjected to antigen retrieval using Antigen Retrieval Solution (Servicebio, G1206) at 90°C for 10 min. Following permeabilization with 0.2% Triton X-100 (Sigma-Aldrich, T8787) in PBS for 30 min, sections were blocked with 5% bovine serum albumin (BSA) at 37°C for 1-hr. Sections were then incubated with primary antibodies overnight at 4°C, followed by incubation with the corresponding fluorophore-conjugated secondary antibodies at 37°C for 1-hr. Nuclei were counterstained with DAPI (Thermo Fisher Scientific, 62248; 1:1,000) for 1 min at room temperature. Slides were mounted using ProLong^®^ Gold Antifade Reagent (Thermo Fisher Scientific, P36930).

For BrdU staining, sections underwent acid denaturation in 1.5 M HCl at 37°C for 30 min following PBST permeabilization. The sections were neutralized with sodium borate buffer at 37°C for 15 min. Subsequent immunostaining steps were performed as described above.

Images were captured using a Leica TCS SP8 confocal microscope. The primary antibodies used were: GFP (Chicken, Abcam, ab13970, 1:1,000), PAX6 (Rabbit, BioLegend, PRB-278P, 1:500), TBR2/EOMES (Rabbit, Abcam, ab23345, 1:500), Ki-67 (Rabbit, Abcam, ab16667, 1:500), SOX2 (Rat, Invitrogen, 53-9811-82, 1:500), BrdU (Rat, Abcam, ab6326, 1:500), TBR1 (Rabbit, Abcam, ab31940, 1:500), SATB2 (Mouse, Abcam, ab51502, 1:200) and HOPX (Rabbit, Proteintech, 11419-1-AP, 1:500). The secondary antibodies used were: Alexa Fluor 488 Goat anti-Chicken (Abcam, ab150169, 1:1,000), Alexa Fluor 555 Donkey anti-Rabbit (Invitrogen, A31572, 1:1,000), Alexa Fluor 647 Donkey anti-Rabbit (Invitrogen, A21245, 1:1,000), Alexa Fluor 555 Donkey anti-Rat (Invitrogen, A48270, 1:1,000), and Alexa Fluor 647 Donkey anti-Mouse (Abcam, ab150107, 1:1,000).

### Quantification of immunofluorescent staining

IUE embryonic brains were collected with at least three biological replicates per time point. For downstream analysis, only treatment-control pairs exhibiting similar electroporation efficiency and comparable brain regions were included. Each brain section was then delineated with a 100 µm wide region using the Leica confocal microscope annotation function for cell counting.

### Luciferase reporter assays

To evaluate the species-specific enhancer activity of the putative *RHOJ* regulatory element, we identified and extracted orthologous genomic sequences from human (GRCh38, chr14:63223858-63225232), chimpanzee (panTro6, chr14:44033891-44035263), gorilla (gorGor6, chr14:43926278-43927656), and macaque (Mmul_10, chr7:125267958-125269327). These sequences were cloned into the pGL3-Promoter vector (Addgene). Additionally, to assess the functional impact of the identified human-specific single nucleotide change (SNC) at chr14:63224141, we generated a mutant construct based on the human sequence, in which the human-specific allele was reverted to the ancestral state.

SK-N-SH cells were seeded in 24-well plates at a density of 1.5 × 10^5^ cells per well. After 12-hr, cells were transfected using Lipofectamine 3000 (Thermo Fisher Scientific, L3000015) and 1,000 ng pGL3-promoter luciferase plasmids (Addgene), 30 ng pRL-TK plasmid (Addgene) was co-transfected to control for transfection efficiency. Experiments were performed with 4 technical replicates (quadruplicate wells) per group.

Luciferase assays were performed 48-hr after transfection using Dual-Luciferase^®^ Reporter Assay System (Promega, E1960) according to the manufacturer’s instructions. The luciferase activity was measured and quantified using TECAN Spark^®^. For each experiment, the raw value was normalized using the pGL3-promoter empty vector. Data were collected from independent differentiations at least three times.

### Generation and validation of *RHOJ* enhancer-deletion cell lines

The H9 human embryonic stem cell (hESC) line was used to generate *RHOJ* enhancer-deletion clones. The hESC line was provided by the Prof. Ming Li’s laboratory of Kunming Institute of Zoology, Chinese Academy of Sciences, in accordance with the agreement with WiCell (WiCell Agreement No., 25-W0392). Two gRNAs were designed to flank a region within the intronic *RHOJ* enhancer containing the human-specific single-nucleotide change tested in the reporter assay. Guide constructs were prepared at 500 ng/µL and introduced into H9 hESCs using Lipofectamine 3000. Solution A was prepared by combining 50 µL Opti-MEM with 5 µL Lipofectamine 3000, and solution B contained 2 µg plasmid DNA diluted in Opti-MEM. Solution B was added to solution A, incubated at room temperature for 10 min and applied to the cells. The medium was replaced after 48 hr. Cells carrying both guide constructs were isolated by flow cytometry and expanded to establish two independently derived enhancer-deletion clones, Enh-KO #1 and Enh-KO #2.

Deletion was assessed by genomic PCR, agarose gel electrophoresis and Sanger sequencing of the deletion junction. TA cloning followed by Sanger sequencing confirmed biallelic deletion in both clones, with alleles carrying 363 bp and 364 bp deletions. The two alleles were supported by seven and three sequenced TA clones in Enh-KO #1 and by three and five TA clones in Enh-KO #2, respectively. The guide-cloning oligonucleotide sequences and genotyping primers are listed below. All sequences are written from 5’ to 3’; lowercase ‘cacc’ denotes the cloning overhang in the supplied guide oligonucleotides.

| Oligonucleotide | Sequence (5’ -3’) |

| RHOJ-Enh-ko sgRNA1 cloning oligonucleotide | caccGTTGATAACATAGCGCTCCAG |

| RHOJ-Enh-ko sgRNA2 cloning oligonucleotide | caccGTGTACTGTAACTCATGCGCA |

| RHOJ-Enh-ko genotyping primer F | TCATTTCATAGTCCAATTTCACTCAAT |

| RHOJ-Enh-ko genotyping primer R | CTTGATTAGGAACGATTCTTCTGGTA |

### Prediction and examination of potential CRISPR off-target sites

Potential off-target sites were identified using Cas-OFFinder (http://www.rgenome.net/cas-offinder/). Genomic sites with one to three base mismatches to either guide, excluding the intended target region, were considered potential off-target sites. Three candidate sites on chromosomes 2, 4 and 8 were examined in WT, Enh-KO #1 and Enh-KO #2 cells. Sequences extending 150 bp upstream and downstream of each candidate site were used to design PCR assays, and the resulting amplicons were analysed by Sanger sequencing. The candidate intervals and primers are listed below.

| Candidate interval | Forward primer (5’ -3’) | Reverse primer (5’ -3’) |

| chr2:161764451-161764751 | GCCACCGCATTTCATTGTTA | CTGCTTTGTCCTCTATTCATTC |

| chr4:41888744-41889044 | GGAAGGTGTGGAAAGGGAAAGG | CTCTTGTGAAGATGAGGCAGCT |

| chr8:36552130-36552430 | GCTGAGGAGGCAGAATTGAAC | GCGGAGAAATTCCTATGTGACA |

### Cortical organoid differentiation

WT H9 hESCs and the two enhancer-deletion clones were differentiated into cortical organoids. On day (D) 0, cells were dissociated into single cells using Accutase (Sigma-Aldrich, A6964) and seeded at 10,000 cells per well in round-bottom, ultra-low-attachment 96 well plates (Corning, 7007), with 150 µL embryoid body (EB) formation medium per well. EB formation medium consisted of DMEM/F12 (Gibco, 31331093), 20% KnockOut Serum Replacement (Thermo Fisher Scientific, 10828028), 1× GlutaMAX (Gibco, 35050061), 1× non-essential amino acids (NEAA; Gibco, 11140050), 1× penicillin/streptomycin (Gibco, 15070063), 0.1% β-mercaptoethanol, 10 µM SB431542 (Selleck, S1067), 10 µM DMH-1 (Tocris, 4126) and 20 µM Y-27632 (Selleck, S1049). On D3, 150 µL fresh EB formation medium was gently added to each well.

On D5, organoids were transferred to the CERO system for three-dimensional suspension culture, and half of the medium was replaced with neural induction medium. Neural induction medium consisted of DMEM/F12 supplemented with 1× GlutaMAX, 1× NEAA, 1× penicillin/streptomycin, 1× N2 (Gibco, 17502048), 1× B27 (Gibco, 17504044) and 2.5 µg/mL heparin (MCE, HY-17567). On D7, the medium was completely replaced with fresh neural induction medium.

On D11, organoids were transferred to differentiation medium consisting of a 1:1 mixture of DMEM/F12 and Neurobasal medium supplemented with 1× GlutaMAX, 1× NEAA, 1× penicillin/streptomycin, 1× N2, 1× B27 and 2.5 µg/mL insulin (MCE, HY-P0035). From D30 onward, organoids were maintained in maturation medium consisting of differentiation medium supplemented with 10 µM forskolin (Selleck, S2449), 0.2 mM ascorbic acid (STEMCELL Technologies, 72132), 10 ng/mL BDNF (PeproTech, 450-02) and 10 ng/mL GDNF (PeproTech, 450-10). Differentiation and maturation media were changed every 5 days.

### Quantification of organoid RNA-FISH

RNAscope images were analyzed using ImageJ. Neural rosette areas were manually outlined as regions of interest (ROIs) based on the organization of DAPI-stained nuclei. Within each ROI, DAPI-positive nuclei were segmented by thresholding and counted using the Analyze Particles function, with a minimum particle size of 12 pixels. *RHOJ* and *HOPX* puncta were detected in their respective channels following background subtraction and thresholding, using identical parameters across experimental groups and a minimum particle size of 4 pixels. For each ROI, the number of *RHOJ* or *HOPX* puncta was divided by the number of DAPI-positive nuclei to obtain the normalized RNA-FISH signal. Multiple neural rosette areas were analyzed for each cell line, with the numbers of organoids and regions reported in the figure legends.

### Quantification of organoid size and immunofluorescence

Organoid diameters were measured at D5, D30, D45, D60 and D75. Cellular analyses were performed in neural rosette areas at D50 and D75. The proportions of HOPX-, SOX2-, TBR1-, TBR2-, Ki67-and SATB2-positive cells were calculated relative to DAPI-positive cells within the analyzed region. The thicknesses of the SOX2-positive progenitor zone and TBR1-positive neuronal layer were measured in D50 organoids. Measurements from Enh-KO #1 and Enh-KO #2 were displayed together as Enh-KO in the cellular quantification panels, with each point representing one neural rosette area. The numbers of organoids and regions analyzed are provided in the figure legends.

### Graphical generation

Schematic elements in Figs. 1, 4, 5 and 6 were created with BioRender.com. The corresponding BioRender records are available at: Fig. 1, https://BioRender.com/nmady8z; Fig. 4, https://BioRender.com/gvrvzo9; Fig. 5, https://BioRender.com/c8bnpur; and Fig. 6, https://BioRender.com/ynv8trx.

### Data Availability

The raw and processed snMultiome data for macaque and mouse developing PFC have been deposited at GEO under accession number: GSE318474.

### Code Availability

Scripts and intermediate data used in this study are available at the GitHub repository: https://github.com/yimingsun12138/CortexDev

## Acknowledgements

We thank Yan Guo for her support and assistance in this study. We also thank Xin Cai and Yue Zhang for their technical advice with cerebral organoid culture. This study was supported by grants from the National Key Research and Development Program of China (2025YFC3410300 to B.S.; 2021YFA1100300 to C.L.), the Major Scientific Project of Yunnan Province of China (202305AH340007 to B.S.), the National Natural Science Foundation of China (32288101 to B.S.; 32288102 and 32025006 to C.L.; 32422018 to X.L.), the Yunnan Scientist Workshop (to B.S.), and the Key Basic Research Project of Yunnan Province of China (202401BC070014 to B.S.; 202501AS070085 to X.L.). Yunnan Revitalization Talent Support Program Top team-202505AT350003 and 202405AS350022 to X.L.).

Part of the data analysis was performed on the High Performance Computing Platform of the Center for Life Sciences, Peking University.

## Author contribution

B.S. conceived and designed the study and supervised the project. C.L. guided and supervised the bioinformatic and computational analyses. Y.S. coordinated the study, performed the bioinformatic, computational and statistical analyses of all multiomic datasets, and wrote the manuscript. M.L. provided resource. K.C. conducted experimental validation with X.Y. and B.Y. drafted the experimental sections of the manuscript. B.S., Xin Luo, F.Z. and K.C. designed the developmental sampling schedule for rhesus macaques and mice. K.C., X. L., F.Z., X.Y., B.Y., X.M., X. L., Y. L. and X.G. performed rhesus macaque and mouse sample collection and surgical procedures. K.C. and X. L. performed 10x snMultiome library preparation. Q.C., X.Q. and T.Y. collected human samples. All authors read and approved the final manuscript.

## Competing interests

The authors declare that they have no competing interests.

---

**Extended Data Fig. 1.**
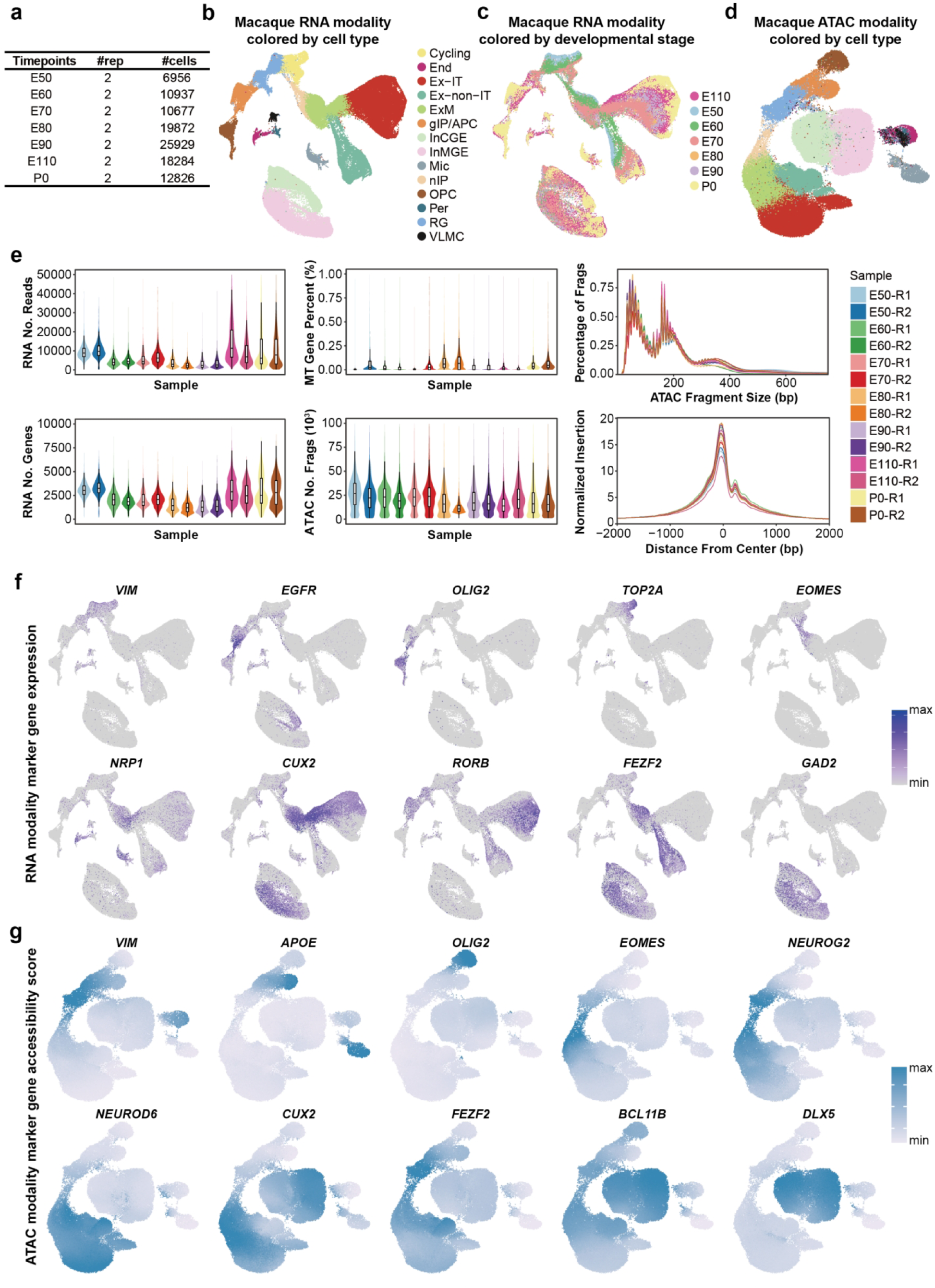
Single-nucleus multiome atlas of the developing macaque cortex. **a**, Developmental stages, source sample numbers and nucleus counts after quality control (QC), comprising 105,481 nuclei across seven stages. **b**,**c**, Transcriptomic UMAPs coloured by cell type (**b**) or developmental stage (**c**). **d**, Chromatin accessibility UMAP coloured by the cell-type annotation of each paired nucleus. **e**, Sample-level QC metrics. Violin plots show RNA counts (upper left), detected genes (lower left), mitochondrial transcript percentage (upper middle) and ATAC fragment counts (lower middle) per nucleus. Curves show ATAC fragment-length distributions (upper right) and normalized Tn5 insertion profiles around transcription start sites (lower right). **f**, Expression of canonical marker genes in the transcriptomic UMAP. **g**, ATAC-derived gene activity scores for canonical markers in the accessibility UMAP. Colour intensity indicates the displayed expression or gene activity score. Cell-type abbreviations are defined in Fig. 1. E, embryonic day; P0, postnatal day 0.

**Extended Data Fig. 2.**
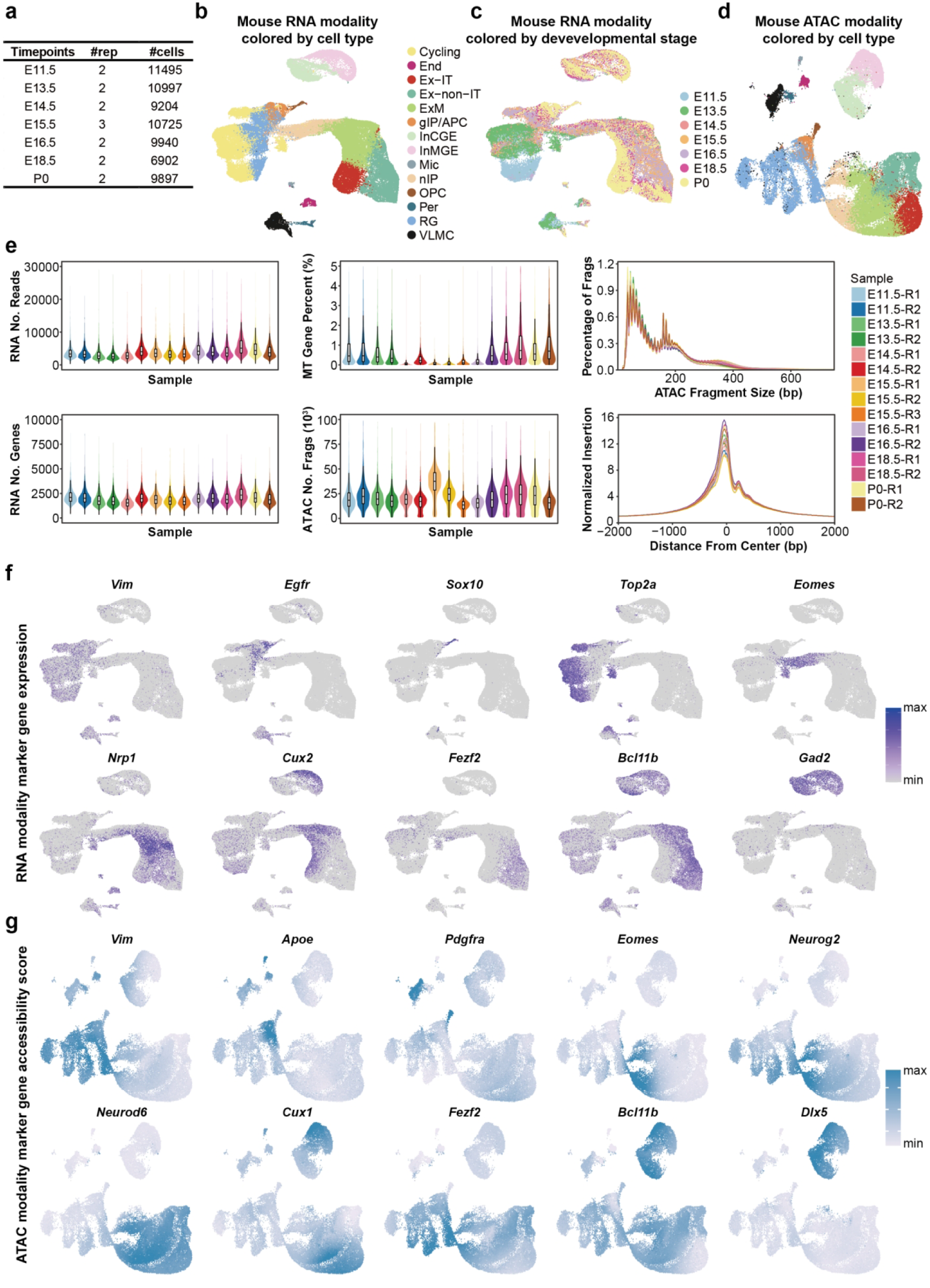
Single-nucleus multiome atlas of the developing mouse cortex. **a**, Developmental stages, source sample numbers and nucleus counts after quality control (QC), comprising 69,160 nuclei across seven stages. **b**,**c**, Transcriptomic UMAPs coloured by cell type (**b**) or developmental stage (**c**). **d**, Chromatin accessibility UMAP coloured by the cell-type annotation of each paired nucleus. **e**, Sample-level QC metrics. Violin plots show RNA counts (upper left), detected genes (lower left), mitochondrial transcript percentage (upper middle) and ATAC fragment counts (lower middle) per nucleus. Curves show ATAC fragment-length distributions (upper right) and normalized Tn5 insertion profiles around transcription start sites (lower right). **f**, Expression of canonical marker genes in the transcriptomic UMAP. **g**, ATAC-derived gene activity scores for canonical markers in the accessibility UMAP. Colour intensity indicates the displayed expression or gene activity score. Cell-type abbreviations are defined in Fig. 1. E, embryonic day; P0, postnatal day 0.

**Extended Data Fig. 3.**
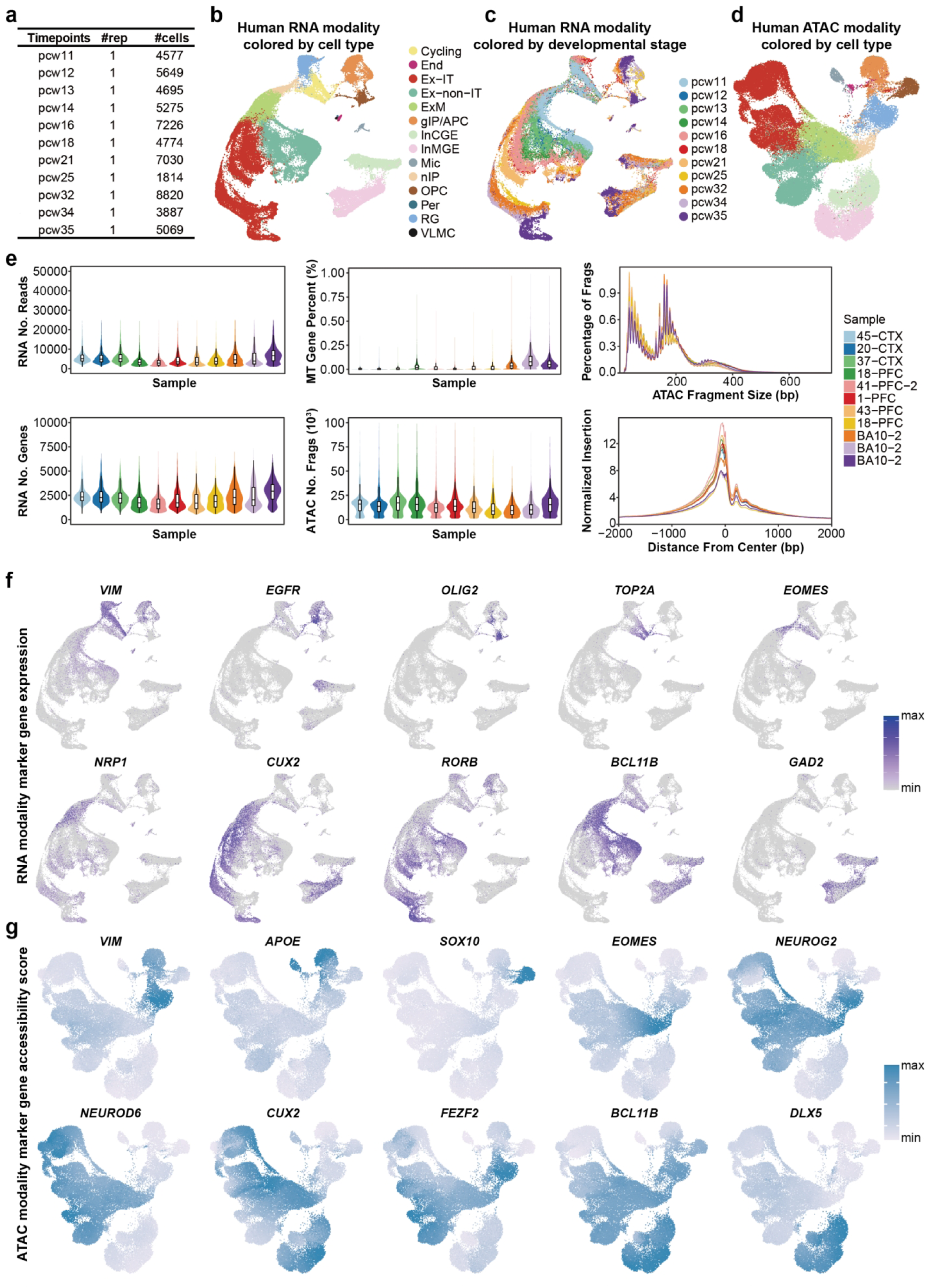
Single-nucleus multiome atlas of the developing human cortex. **a**, Developmental stages, source sample numbers and nucleus counts after quality control (QC) from the published dataset of Wang et al. (2025), comprising 58,816 nuclei across 11 stages. **b,c**, Transcriptomic UMAPs coloured by cell type (b) or developmental stage (c). **d**, Chromatin accessibility UMAP coloured by the cell-type annotation of each paired nucleus. **e**, Sample-level QC metrics. Violin plots show RNA counts (upper left), detected genes (lower left), mitochondrial transcript percentage (upper middle) and ATAC fragment counts (lower middle) per nucleus. Curves show ATAC fragment-length distributions (upper right) and normalized Tn5 insertion profiles around transcription start sites (lower right). **f**, Expression of canonical marker genes in the transcriptomic UMAP. **g**, ATAC-derived gene activity scores for canonical markers in the accessibility UMAP. Colour intensity indicates the displayed expression or gene activity score. Cell-type abbreviations are defined in Fig. 1. pcw, post-conception week.

**Extended Data Fig. 4.**
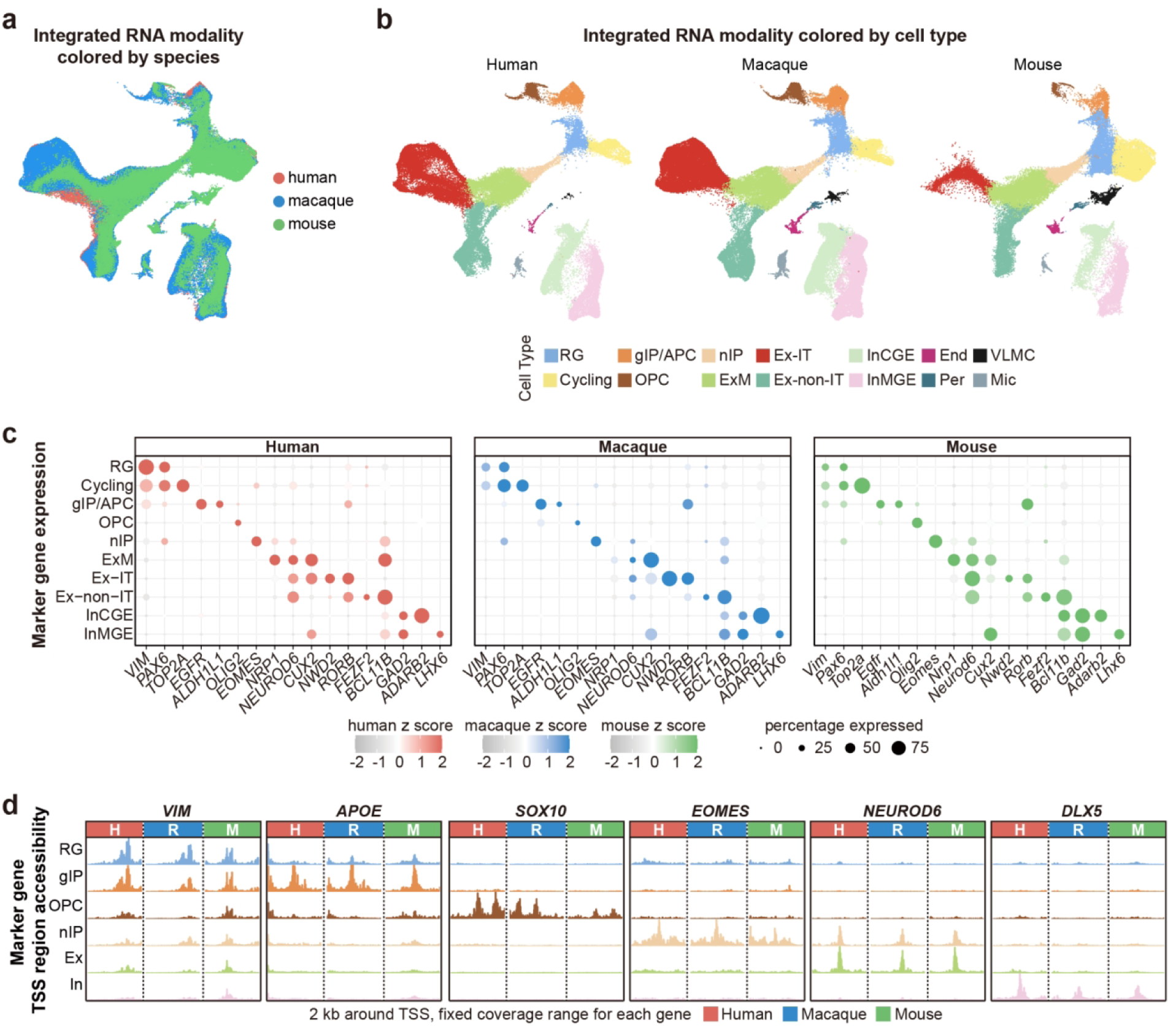
Cross-species integration and cortical cell-type characterization. **a**, Integrated human, macaque and mouse transcriptomic UMAP coloured by species. **b**, The same embedding shown separately for each species and coloured by cell type. **c**, Canonical marker expression across cell types. Dot size indicates the percentage of nuclei expressing the gene; colour indicates the gene-wise expression z score within each species. **d**, Aggregated accessibility profiles around canonical marker transcription start sites (TSSs). Signals for each gene are displayed on a fixed vertical scale across cell types and species. H, human; R, rhesus macaque; M, mouse; Ex, excitatory neurons; In, inhibitory neurons. The gIP label denotes gIP/APC. Other cell-type abbreviations are defined in Fig. 1.

**Extended Data Fig. 5.**
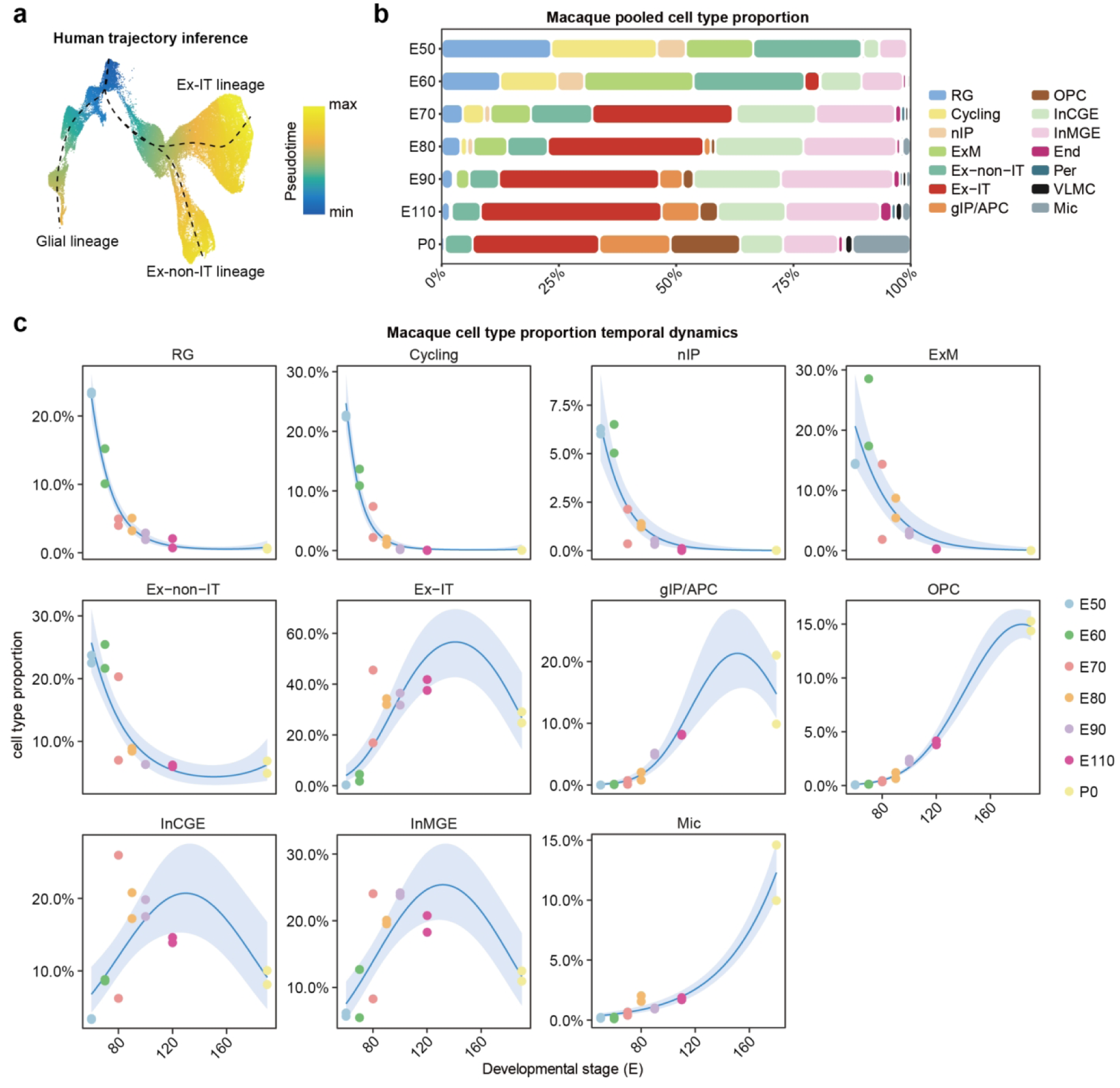
Inferred developmental trajectories and cell-type composition in the macaque cortex. **a**, UMAP of cortical progenitor, excitatory neuronal and glial populations coloured by inferred pseudotime. Dashed lines show Slingshot-inferred trajectories rooted in RG and extending towards Ex-IT, Ex-non-IT and glial populations. **b**, Cell-type proportions at each developmental stage, calculated from nuclei pooled across samples at that stage. **c**, Sample-level developmental profiles of cell-type proportions. The denominator is all retained annotated nuclei in each sample. Each point represents a source sample (n = 14 samples), coloured by stage. Solid lines show beta-binomial regression fits; shaded bands indicate 95% confidence intervals for the fitted mean. P0 was assigned E180 for continuous-age fitting. Cell-type abbreviations are defined in Fig. 1. E, embryonic day; P0, postnatal day 0.

**Extended Data Fig. 6.**
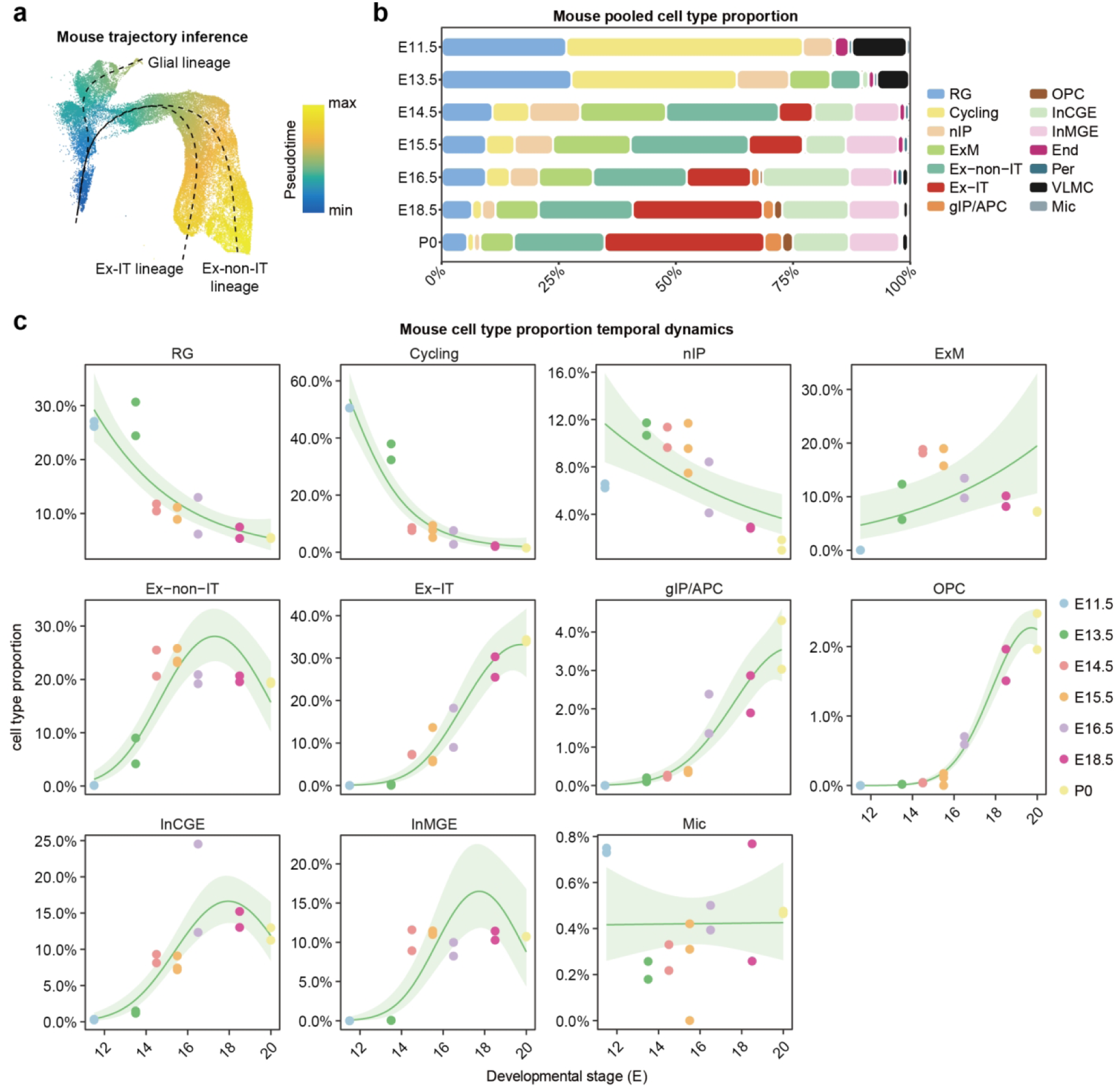
Inferred developmental trajectories and cell-type composition in the mouse cortex. **a**, UMAP of cortical progenitor, excitatory neuronal and glial populations coloured by inferred pseudotime. Dashed lines show Slingshot-inferred trajectories rooted in RG and extending towards Ex-IT, Ex-non-IT and glial populations. **b**, Cell-type proportions at each developmental stage, calculated from nuclei pooled across samples at that stage. **c**, Sample-level developmental profiles of cell-type proportions. The denominator is all retained annotated nuclei in each sample. Each point represents a source sample (n = 15 samples), coloured by stage. Solid lines show beta-binomial regression fits; shaded bands indicate 95% confidence intervals for the fitted mean. P0 was assigned E20 for continuous-age fitting. Cell-type abbreviations are defined in Fig. 1. E, embryonic day; P0, postnatal day 0.

**Extended Data Fig. 7.**
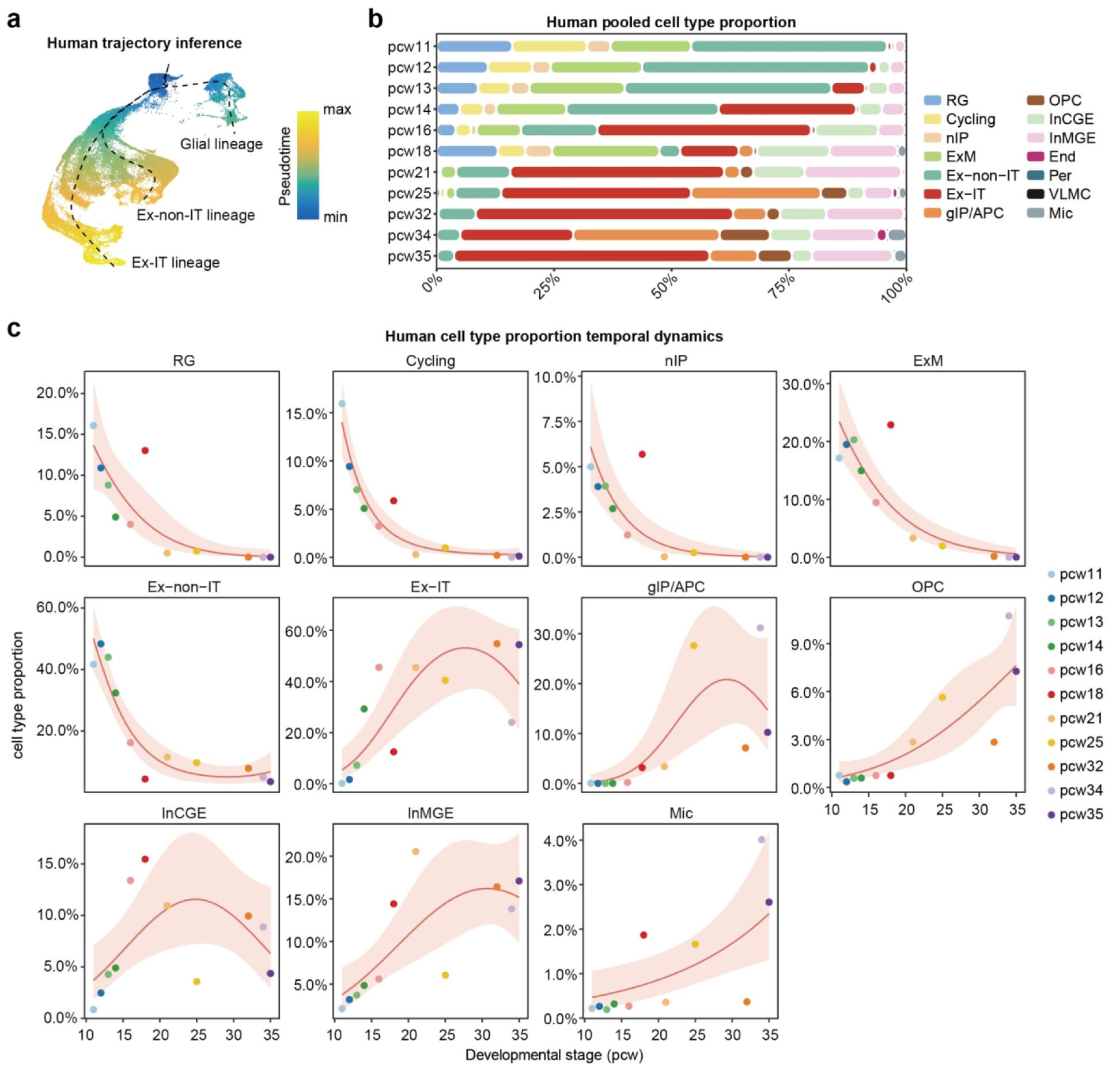
Inferred developmental trajectories and cell-type composition in the human cortex. **a**, UMAP of cortical progenitor, excitatory neuronal and glial populations coloured by inferred pseudotime. Dashed lines show Slingshot-inferred trajectories rooted in RG and extending towards Ex-IT, Ex-non-IT and glial populations. **b**, Cell-type proportions at each developmental stage, calculated from nuclei pooled across samples at that stage. **c**, Sample-level developmental profiles of cell-type proportions. The denominator is all retained annotated nuclei in each sample. Each point represents a source sample (n = 11 samples), coloured by stage. Solid lines show beta-binomial regression fits; shaded bands indicate 95% confidence intervals for the fitted mean. Cell-type abbreviations are defined in Fig. 1. pcw, post-conception week.

**Extended Data Fig. 8.**
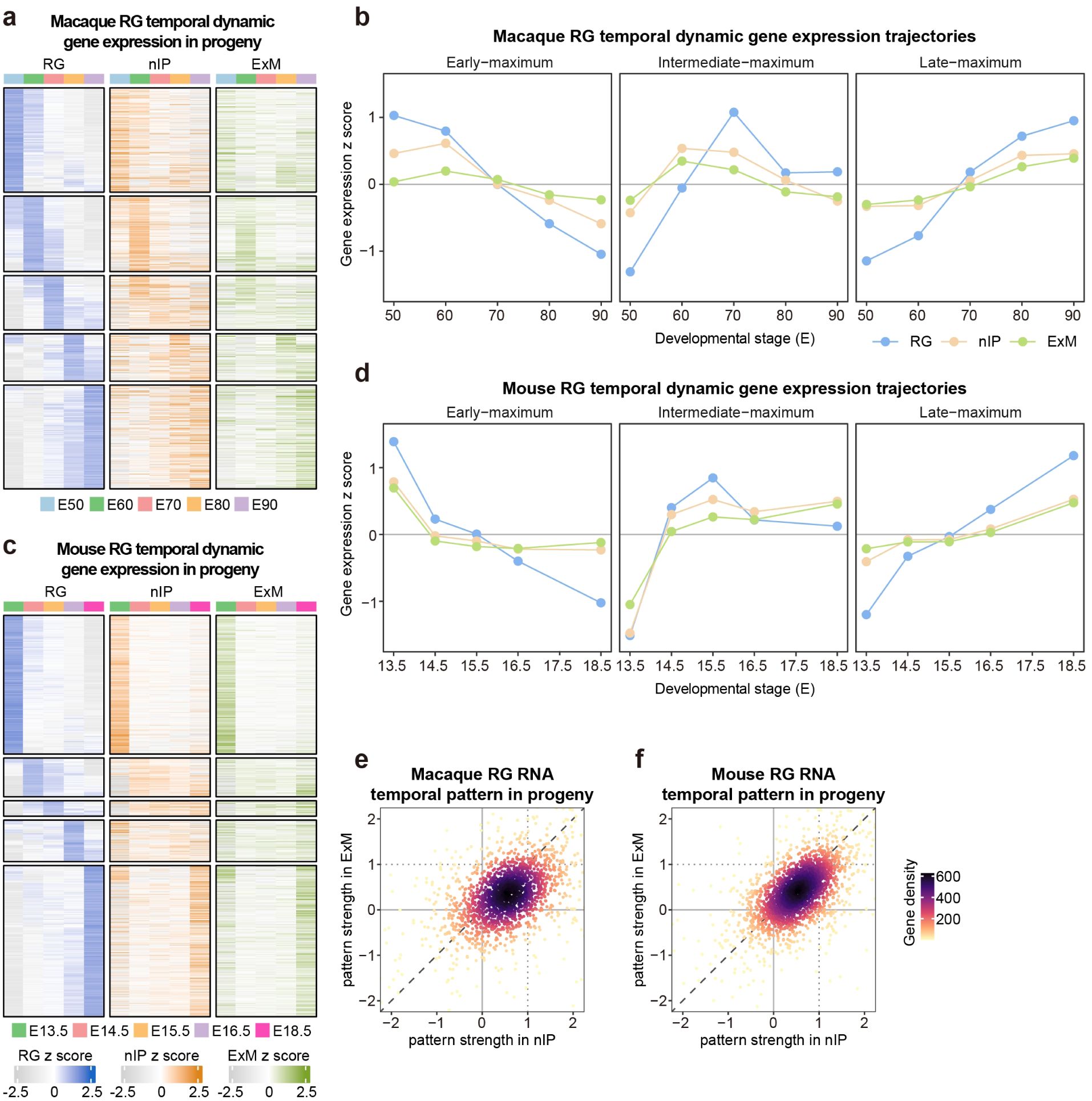
Persistence of RG temporal expression signatures in macaque and mouse. **a**,**c**, Expression profiles of temporally dynamic RG genes in RG, nIP and ExM from macaque (**a**; 1,979 genes, E50-E90) and mouse (**c**; 2,650 genes, E13.5-E18.5). Genes are grouped and ordered by their stage of maximum RG expression. The RG-defined row order and numerical scale are retained across all three cell types. **b**,**d**, Median profiles of genes with early, intermediate or late RG expression maxima in macaque (**b**) and mouse (**d**), using the same RG-referenced scale. Early and late groups comprise genes peaking at the first two and last two displayed stages, respectively; the intermediate group comprises genes peaking at the middle stage. Points show group medians, connected by lines and coloured by cell type. **e**,**f**, Relative RG temporal pattern strength in nIP and ExM for macaque (**e**) and mouse (**f**), calculated as in Fig. 1g. Each point represents a gene; colour indicates local gene density. Dashed diagonals indicate equal strength in nIP and ExM; dotted lines indicate the RG reference strength of 1. RG, radial glia; nIP, neuronal intermediate progenitors; ExM, migrating excitatory neurons; E, embryonic day.

**Extended Data Fig. 9.**
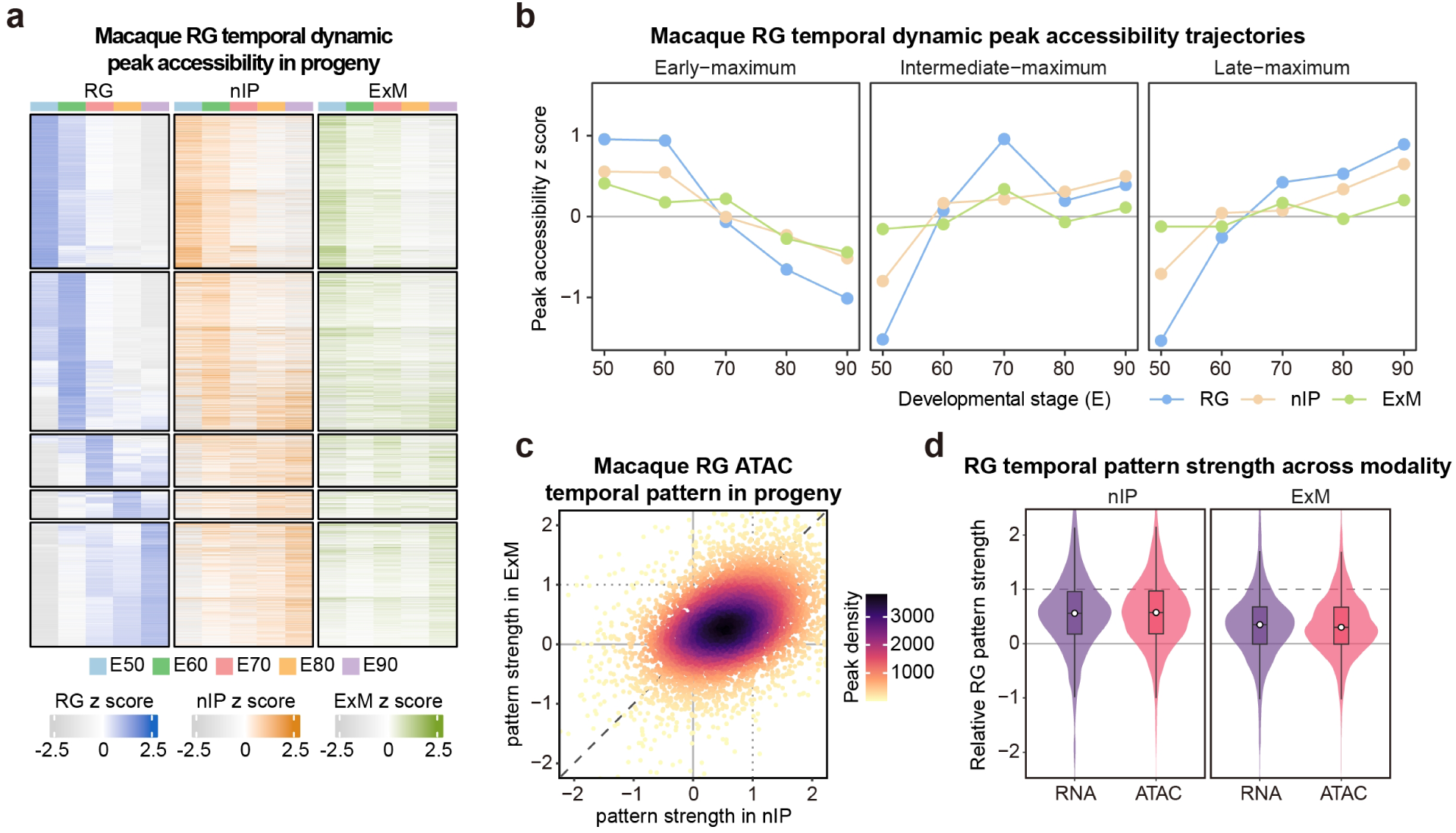
Persistence of RG temporal accessibility signatures in macaque nIP and ExM. **a**, Accessibility profiles of 9,070 temporally dynamic RG peaks in RG, nIP and ExM across E50-E90. Peaks are grouped and ordered by their stage of maximum RG accessibility. The RG-defined row order and numerical scale are retained across cell types. **b**, Median accessibility profiles of peaks with early (E50-E60), intermediate (E70) or late (E80-E90) RG maxima. Points show group medians, connected by lines and coloured by cell type. **c**, Relative RG temporal accessibility pattern strength in nIP and ExM, calculated as the regression slope of the downstream profile against the corresponding RG profile across matched stages. Each point represents a peak; colour indicates local peak density. The dashed diagonal indicates equal strength in nIP and ExM; dotted lines indicate the RG reference strength of 1. **d**, Distributions of relative RG temporal pattern strength for RNA and ATAC in nIP and ExM (1,979 genes and 9,070 peaks). Dashed lines indicate the RG reference strength of 1. RG, radial glia; nIP, neuronal intermediate progenitors; ExM, migrating excitatory neurons; E, embryonic day.

**Extended Data Fig. 10.**
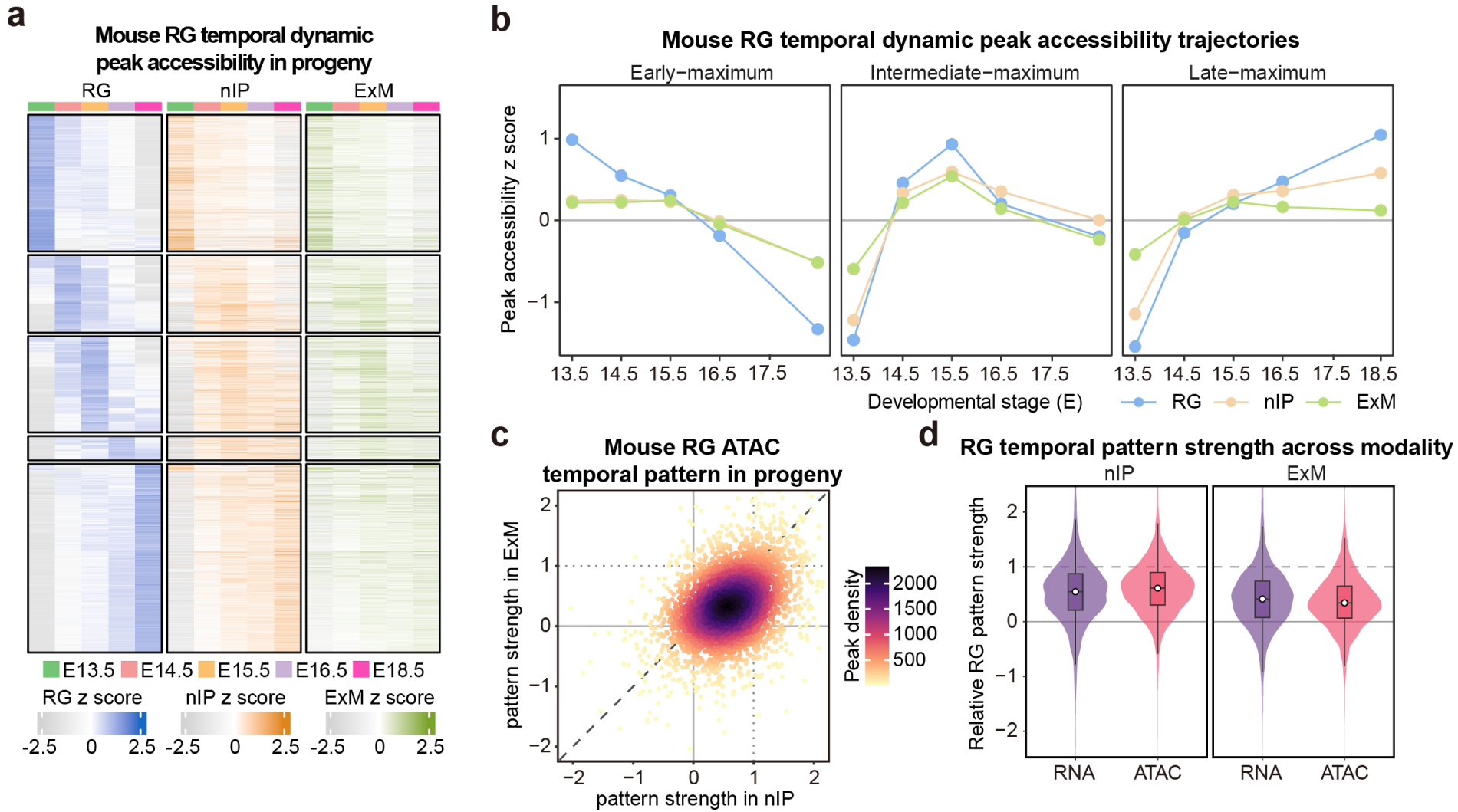
Persistence of RG temporal accessibility signatures in mouse nIP and ExM. **a**, Accessibility profiles of 4,984 temporally dynamic RG peaks in RG, nIP and ExM across E13.5-E18.5. Peaks are grouped and ordered by their stage of maximum RG accessibility. The RG-defined row order and numerical scale are retained across cell types. **b**, Median accessibility profiles of peaks with early (E13.5-E14.5), intermediate (E15.5) or late (E16.5-E18.5) RG maxima. Points show group medians, connected by lines and coloured by cell type. **c**, Relative RG temporal accessibility pattern strength in nIP and ExM, calculated as the regression slope of the downstream profile against the corresponding RG profile across matched stages. Each point represents a peak; colour indicates local peak density. The dashed diagonal indicates equal strength in nIP and ExM; dotted lines indicate the RG reference strength of 1. **d**, Distributions of relative RG temporal pattern strength for RNA and ATAC in nIP and ExM (2,650 genes and 4,984 peaks). Dashed lines indicate the RG reference strength of 1. RG, radial glia; nIP, neuronal intermediate progenitors; ExM, migrating excitatory neurons; E, embryonic day.

**Extended Data Fig. 11.**
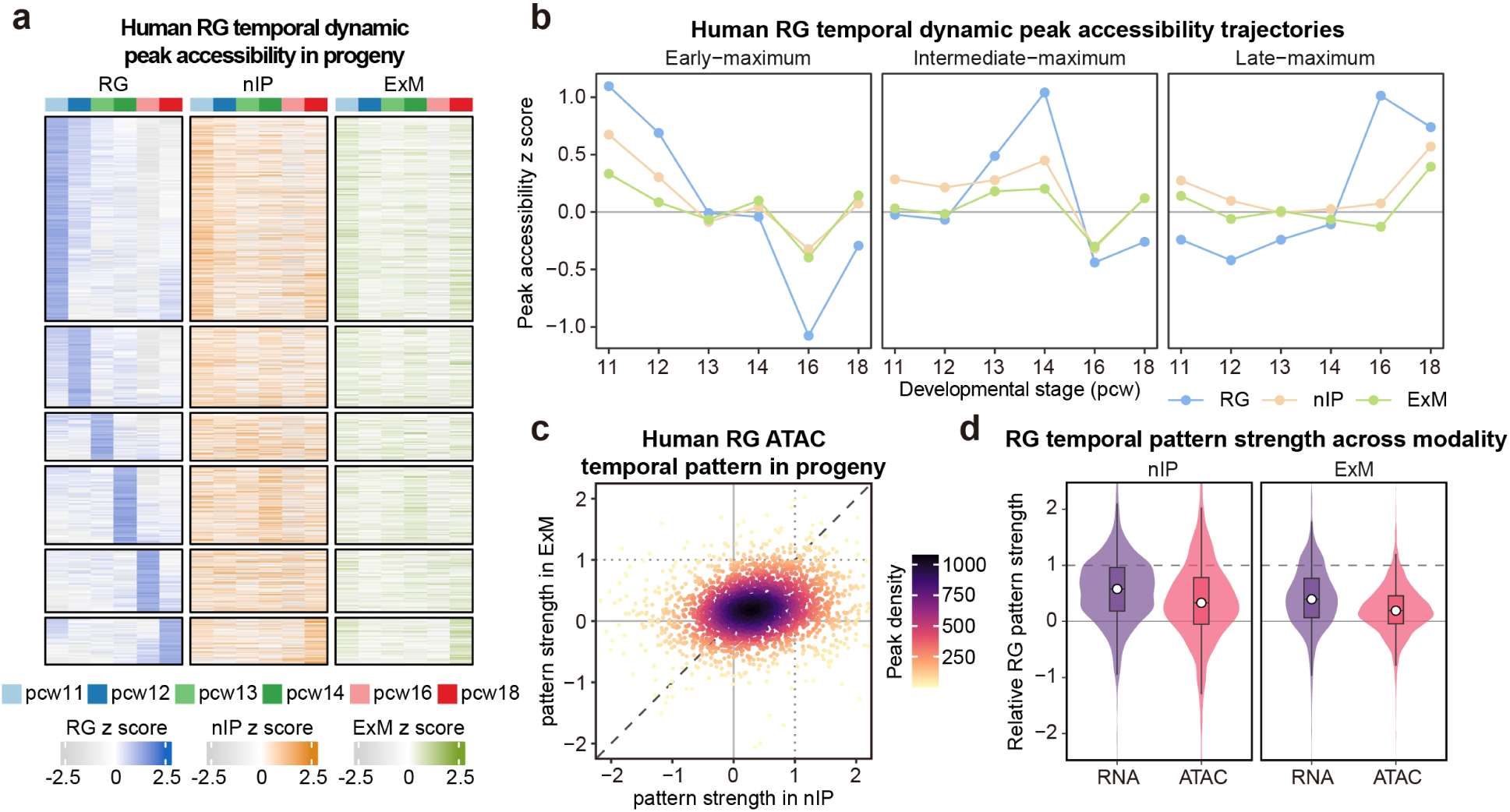
Persistence of RG temporal accessibility signatures in human nIP and ExM. **a,** Accessibility profiles of 2,744 temporally dynamic RG peaks in RG, nIP and ExM across pcw11-18. Peaks are grouped and ordered by their stage of maximum RG accessibility. The RG-defined row order and numerical scale are retained across cell types. **b**, Median accessibility profiles of peaks with early (pcw11-12), intermediate (pcw13-14) or late (pcw16-18) RG maxima. Points show group medians, connected by lines and coloured by cell type. **c**, Relative RG temporal accessibility pattern strength in nIP and ExM, calculated as the regression slope of the downstream profile against the corresponding RG profile across matched stages. Each point represents a peak; colour indicates local peak density. The dashed diagonal indicates equal strength in nIP and ExM; dotted lines indicate the RG reference strength of 1. **d**, Distributions of relative RG temporal pattern strength for RNA and ATAC in nIP and ExM (1,741 genes and 2,744 peaks). Dashed lines indicate the RG reference strength of 1. RG, radial glia; nIP, neuronal intermediate progenitors; ExM, migrating excitatory neurons; pcw, post-conception week.

**Extended Data Fig. 12.**
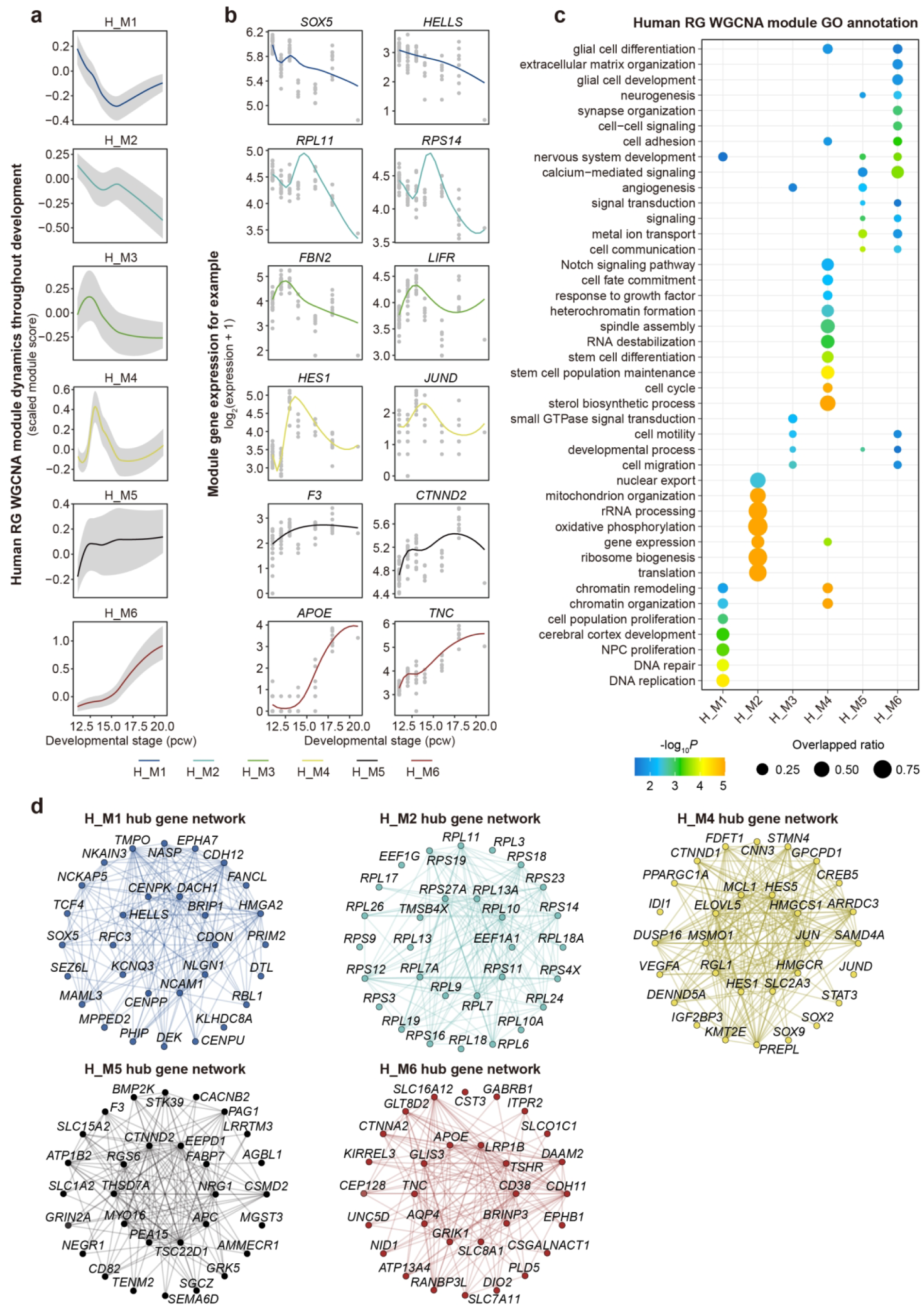
Temporal and functional characterization of human RG modules. **a**, Developmental profiles of six human RG WGCNA modules. Cellular scores are kME-weighted means of gene-wise standardized expression. Coloured lines show LOESS fits; grey bands show separately smoothed stage-wise 25th and 75th percentiles of cellular scores. **b**, Expression profiles of two representative genes per module. Grey points represent RG metacells; coloured lines show generalized additive model (GAM) fits. **c**, Enriched GO biological process terms. Dot size indicates the gene overlap ratio; colour indicates −log₁₀ *P*. **d**, Co-expression networks of 30 hub genes for H_M1, H_M2, H_M4, H_M5 and H_M6. The H_M3 network is shown in Fig. 2c. RG, radial glia; WGCNA, weighted gene co-expression network analysis; kME, module membership; GO, Gene Ontology; pcw, post-conception week.

**Extended Data Fig. 13.**
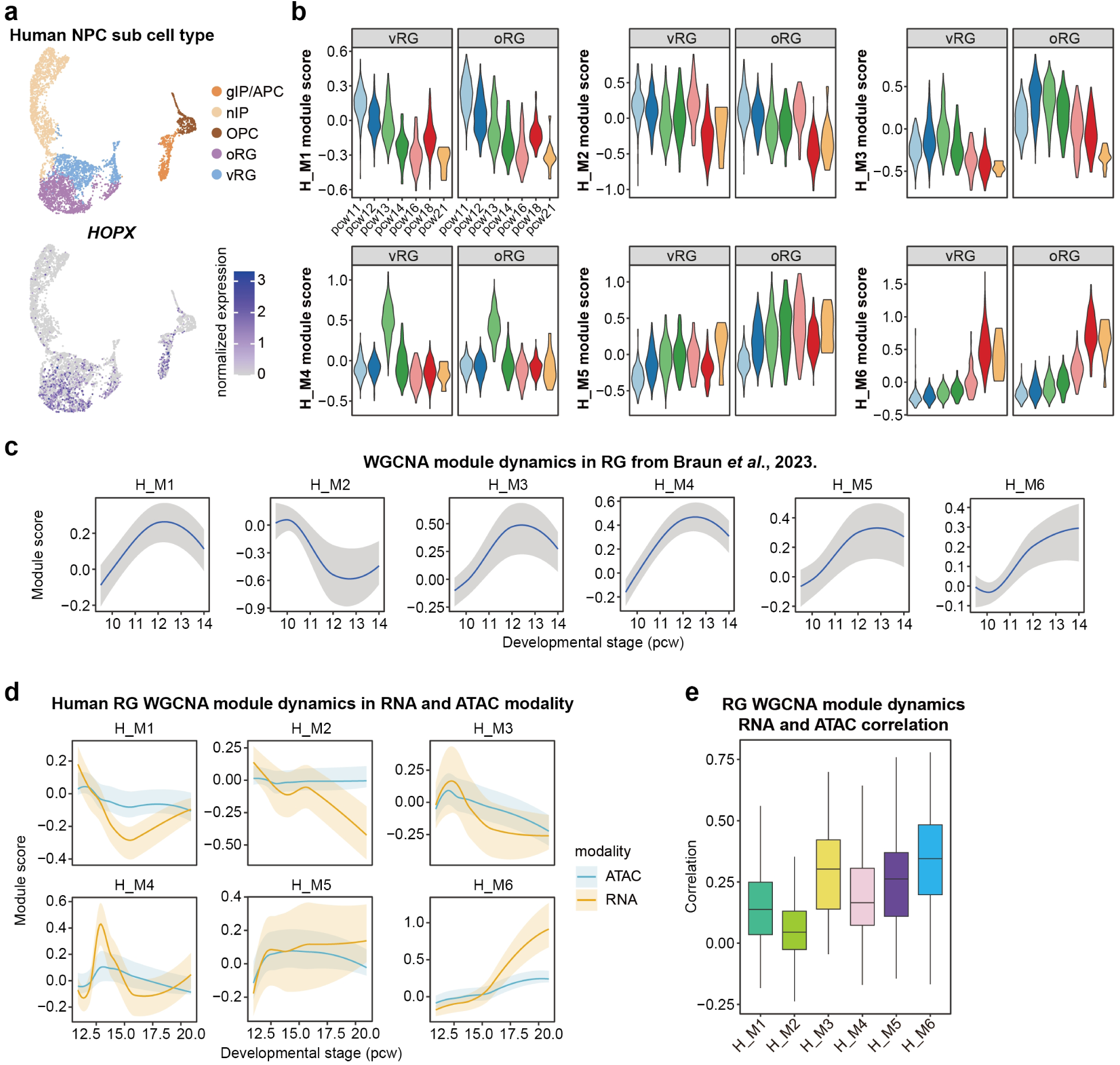
Human RG module profiles across progenitor subtypes, datasets and modalities. **a**, Human neural progenitor UMAP coloured by cell type (top) or normalized *HOPX* expression (bottom). **b**, Cellular module-score distributions in vRG and oRG at each stage. Scores are kME-weighted means of expression standardized gene-wise across the combined vRG and oRG populations. Colours indicate stages. **c**, Module profiles in an independent human cortical dataset (Braun *et al*., 2023), using RG from pcw9.5, pcw10, pcw12 and pcw14. Scores were calculated from gene-wise standardized expression using human kME weights. Lines show LOESS fits; grey bands show separately smoothed stage-wise 25th and 75th percentiles of cellular scores. **d**, Module profiles in RNA expression and ATAC-derived gene activity. Each modality was standardized separately and scored using the same module genes and human kME weights. Lines show LOESS fits; shaded bands show separately smoothed stage-wise 25th and 75th percentiles. **e**, Distributions of gene-wise Pearson correlations between RNA expression and ATAC-derived gene activity across human RG metacells, grouped by module. RG, radial glia; vRG, ventricular radial glia; oRG, outer radial glia; nIP, neuronal intermediate progenitors; gIP/APC, glial intermediate progenitors/astrocyte precursor cells; OPC, oligodendrocyte precursor cells; pcw, post-conception week.

**Extended Data Fig. 14.**
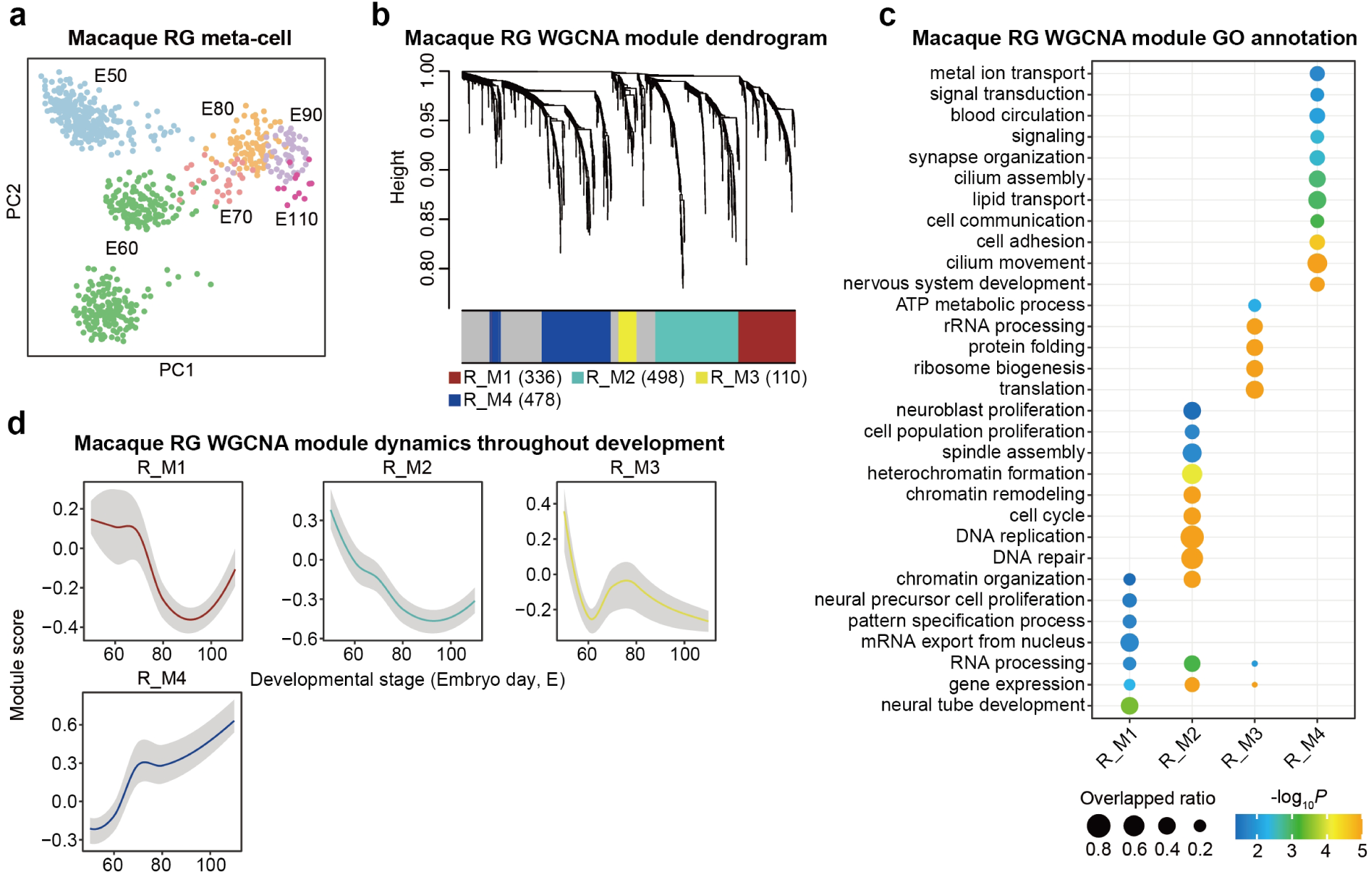
Temporal and functional characterization of macaque RG modules. **a**, Principal component analysis (PCA) of RG metacells coloured by developmental stage. Each point represents a metacell (n = 778). **b**, Hierarchical clustering of temporally dynamic RG genes by WGCNA, resolving four modules. The colour bar indicates module assignments; grey denotes unassigned genes. Numbers in parentheses indicate module sizes. **c**, Enriched GO biological process terms. Dot size indicates the gene overlap ratio; colour indicates −log₁₀ *P*. **d**, Developmental module-score profiles. Scores are kME-weighted means of expression standardized gene-wise within macaque RG. Coloured lines show LOESS fits; grey bands show separately smoothed stage-wise 25th and 75th percentiles of cellular scores. RG, radial glia; WGCNA, weighted gene co-expression network analysis; GO, Gene Ontology; kME, module membership; E, embryonic day.

**Extended Data Fig. 15.**
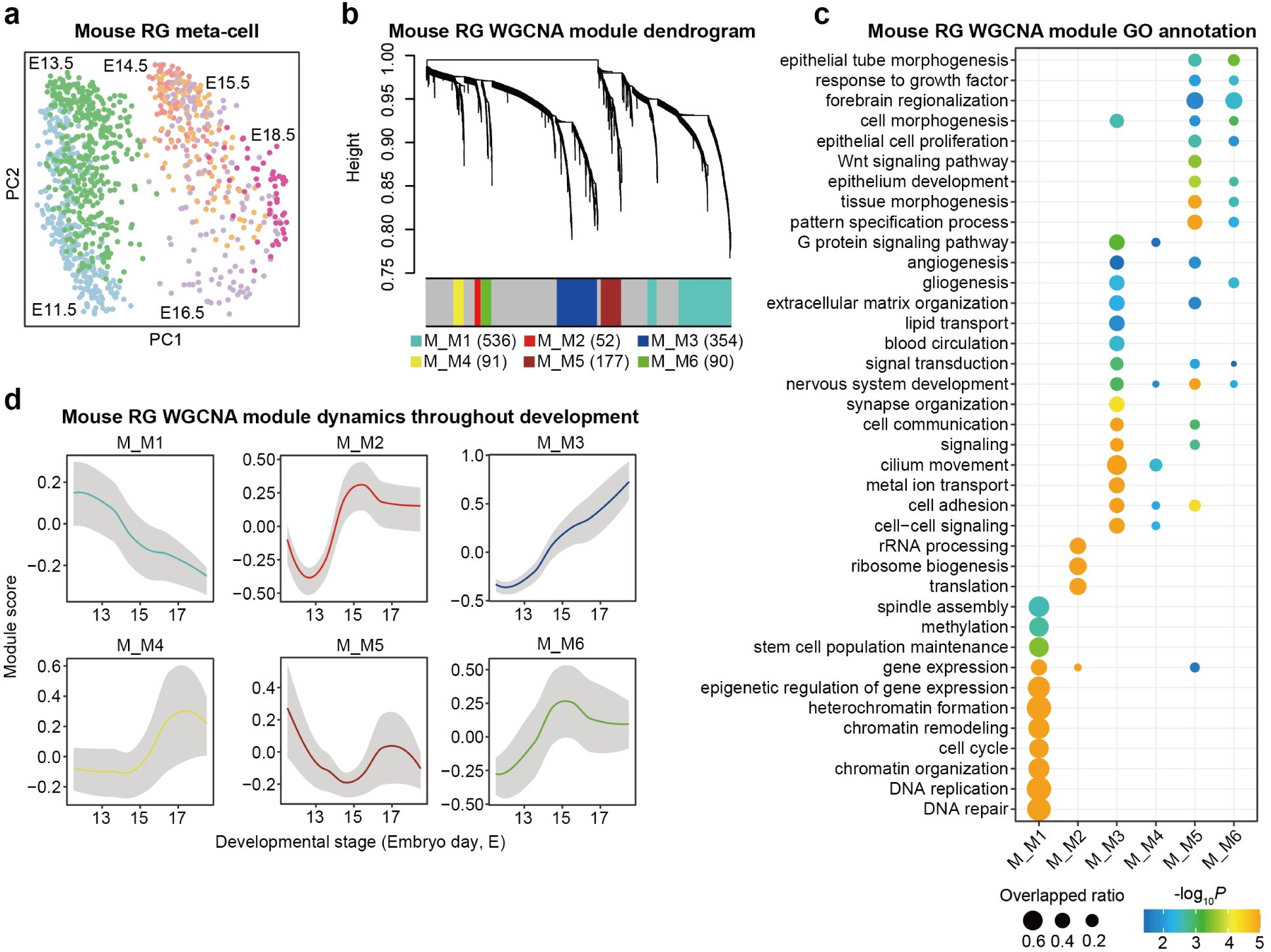
Temporal and functional characterization of mouse RG modules. **a**, Principal component analysis (PCA) of RG metacells coloured by developmental stage. Each point represents a metacell (n = 1,078). **b**, Hierarchical clustering of temporally dynamic RG genes by WGCNA, resolving six modules. The colour bar indicates module assignments; grey denotes unassigned genes. Numbers in parentheses indicate module sizes. **c**, Enriched GO biological process terms. Dot size indicates the gene overlap ratio; colour indicates −log₁₀ P. **d**, Developmental module-score profiles. Scores are kME-weighted means of expression standardized gene-wise within mouse RG. Coloured lines show LOESS fits; grey bands show separately smoothed stage-wise 25th and 75th percentiles of cellular scores. RG, radial glia; WGCNA, weighted gene co-expression network analysis; GO, Gene Ontology; kME, module membership; E, embryonic day.

**Extended Data Fig. 16.**
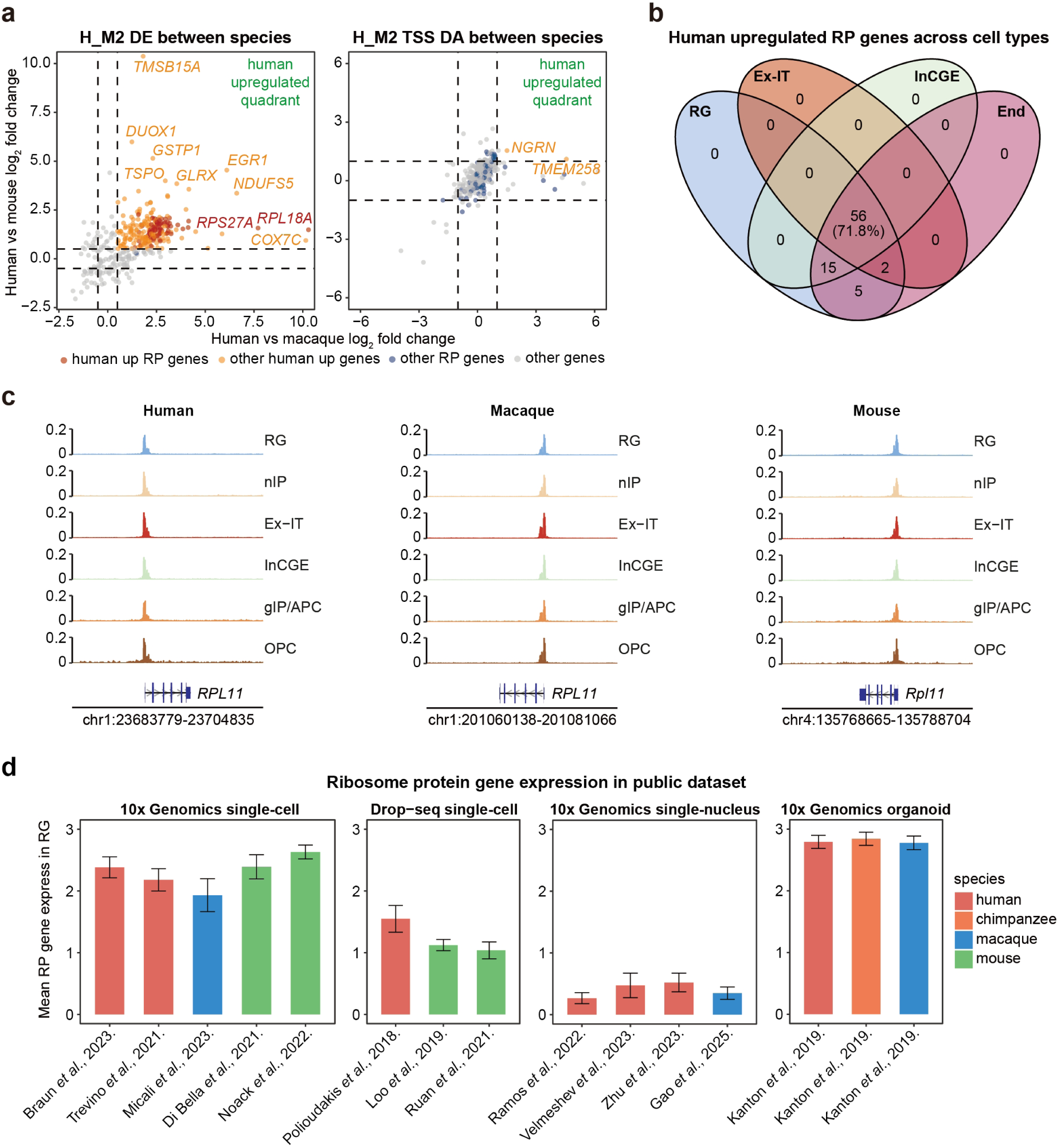
Cross-species expression of ribosome biogenesis-associated genes. **a**, Cross-species differences in H_M2 RNA expression (left) and TSS accessibility (right) in RG. Each point represents a gene; axes show human-to-macaque and human-to-mouse log₂ fold changes. Human-upregulated expression required log₂ fold change > 0.5 and adjusted *P* < 0.01 in both comparisons, using Wilcoxon rank-sum tests. Accessibility was quantified within ±2 kb of TSSs using a ReadsInTSS-normalized peak matrix; human-enriched accessibility required log₂ fold change > 1 and adjusted *P* < 0.01 in both comparisons. Dashed lines indicate fold-change thresholds. Colours distinguish human-upregulated ribosomal protein (RP) genes, other human-upregulated genes, other RP genes and remaining genes. **b**, Overlap of human-upregulated RP genes in RG, Ex-IT, InCGE and endothelial cells, using the RNA criteria in **a**. Numbers indicate genes in each intersection; 56 are shared across all four populations. **c**, Accessibility tracks at the *RPL11* locus in six cell populations across species. Signals are ReadsInTSS-normalized and displayed on a common vertical scale. **d**, Mean RP gene expression in RG from published datasets, grouped by library preparation platform. Dataset identities are shown on the x axes; colours indicate species. Data are mean ± s.d. RG, radial glia; TSS, transcription start site; Ex-IT, intratelencephalic-projecting excitatory neurons; InCGE, caudal ganglionic eminence-derived inhibitory neurons; End, endothelial cells; nIP, neuronal intermediate progenitors; gIP/APC, glial intermediate progenitors/astrocyte precursor cells; OPC, oligodendrocyte precursor cells.

**Extended Data Fig. 17.**
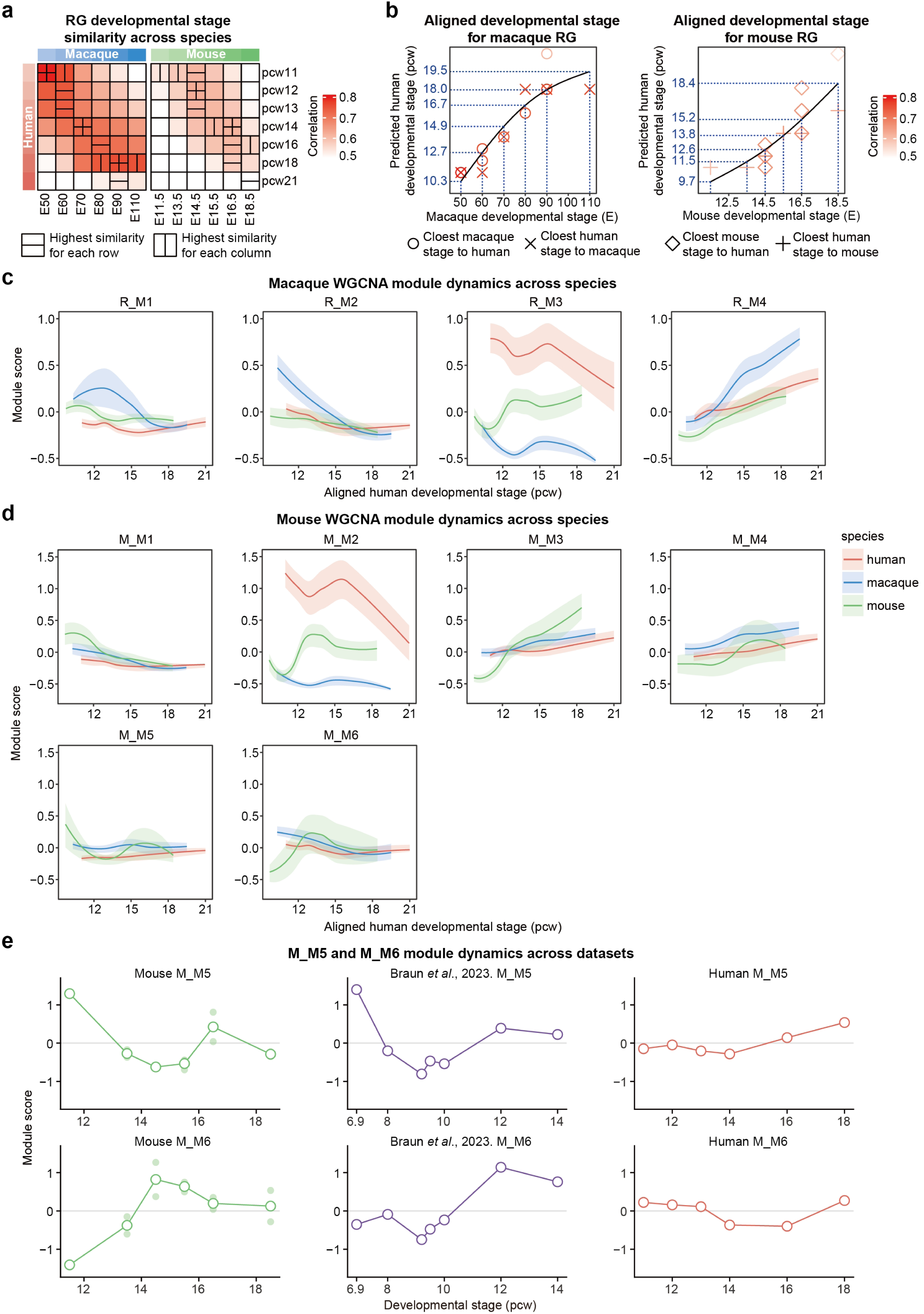
RG developmental stage alignment and cross-species module profiles. **a**, Transcriptomic similarity of human RG stages to macaque (left) and mouse (right) stages. Pearson correlations were calculated between stage-level pseudobulk profiles of shared genes from H_M1, R_M1, R_M2, M_M1, H_M5, H_M6, R_M4, M_M3 and M_M4. Horizontal and vertical marks indicate the maximum correlation within each row and column, respectively. **b**, Alignment of macaque and mouse stages to a human RG developmental timeline. Symbols indicate stage pairs selected by maximum similarity in each direction and are coloured by correlation. Black curves show GAM fits; blue dotted lines indicate assigned human-equivalent ages. **c**,**d**, Macaque (**c**) and mouse (**d**) module profiles across all three species along the aligned timeline. Scores are kME-weighted means of expression standardized gene-wise across the combined species. Lines show LOESS fits; shaded bands show separately smoothed stage-wise 25th and 75th percentiles of cellular scores. Human, red; macaque, blue; mouse, green. **e**, M_M5 and M_M6 profiles in mouse RG, the earlier human dataset of Braun *et al*. (2023), and the human dataset used in this study, plotted against native developmental stages. Open circles show stage-level scores, connected by lines; faint filled points indicate individual samples. RG, radial glia; GAM, generalized additive model; kME, module membership; E, embryonic day; pcw, post-conception week.

**Extended Data Fig. 18.**
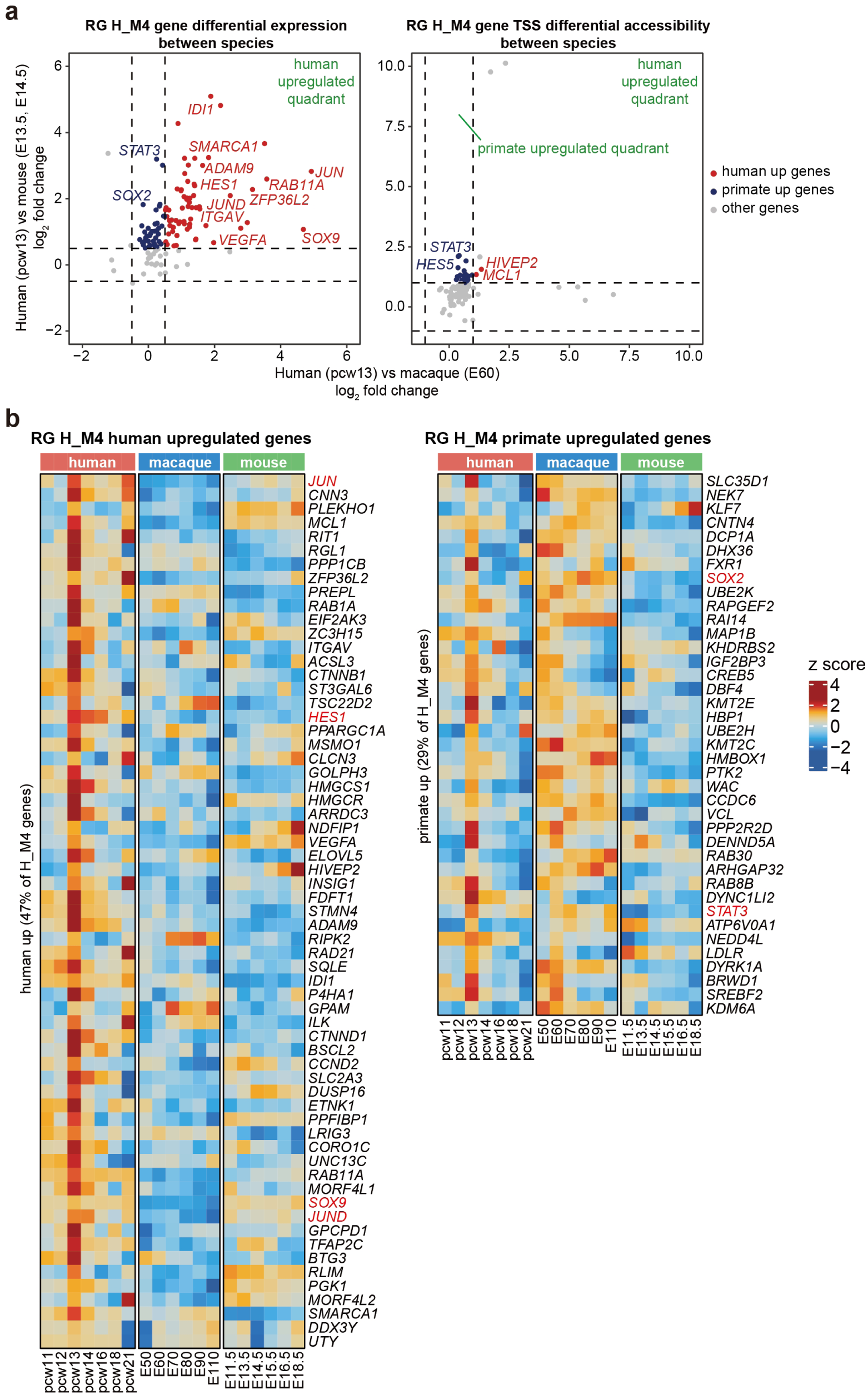
Cross-species expression and TSS accessibility of H_M4 genes. **a**, H_M4 gene expression (left) and TSS accessibility (right) differences between human pcw13 and macaque E60 (x axis), and between human pcw13 and mouse E13.5-14.5 (y axis). Each point represents a gene. Human-upregulated expression required log₂ fold change > 0.5 and adjusted *P* < 0.01 in both comparisons, using Wilcoxon rank-sum tests. Accessibility was measured within ±2 kb of TSSs using a ReadsInTSS-normalized peak matrix; human-enriched accessibility required log₂ fold change > 1 and adjusted *P* < 0.01 in both comparisons. Dashed lines indicate fold-change thresholds. Red, human-upregulated; blue, primate-upregulated; grey, other genes, classified separately for RNA and accessibility. **b**, Temporal expression of human-upregulated (left; 47% of H_M4 genes) and primate-upregulated (right; 29%) genes in RG. Colours show gene-wise z scores across the displayed species and stages. Selected genes are labelled in red. RG, radial glia; TSS, transcription start site; pcw, post-conception week; E, embryonic day.

**Extended Data Fig. 19.**
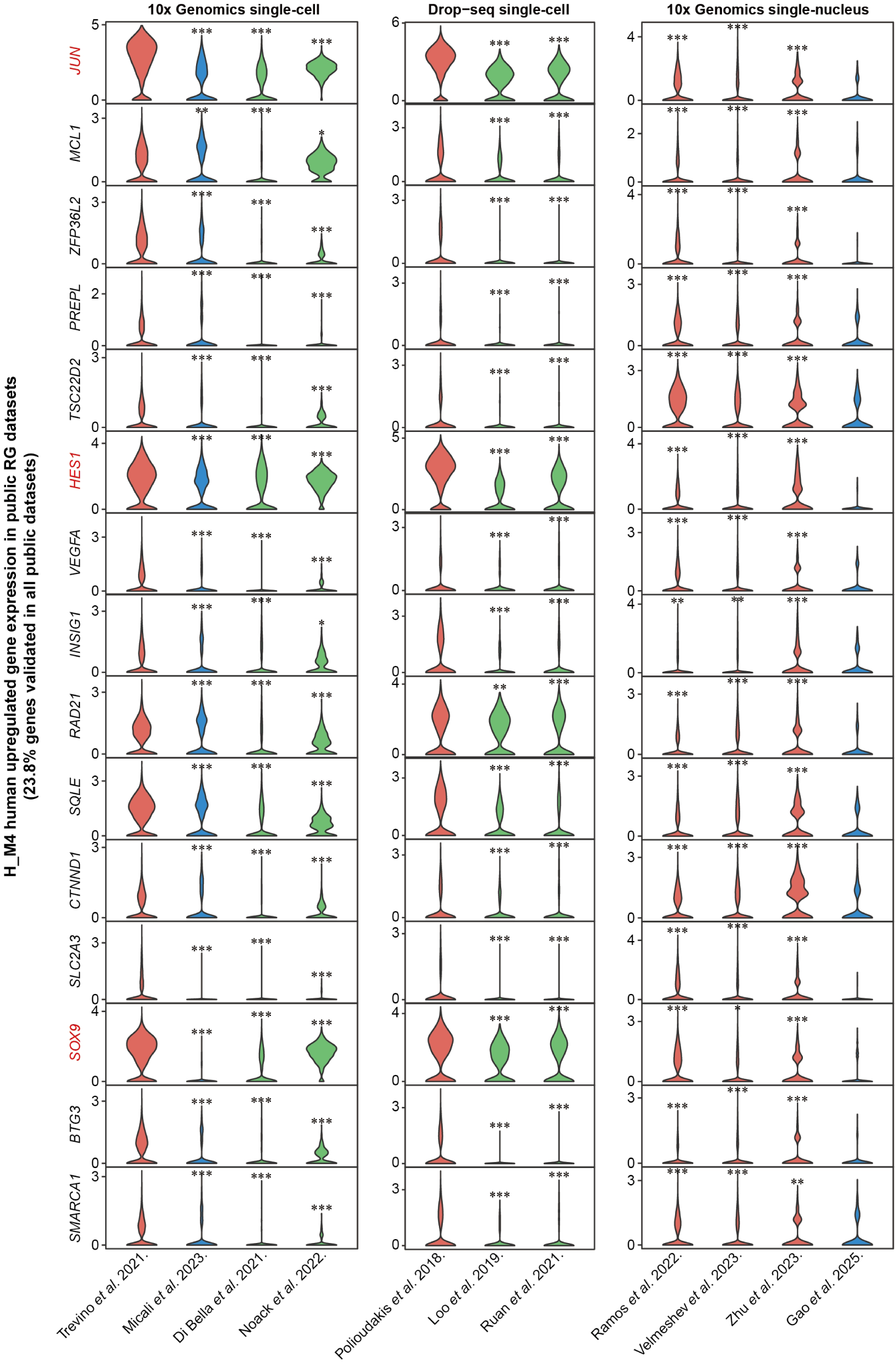
Independent support for human-enriched expression of H_M4 genes. Normalized expression of 15 H_M4 genes in RG from independent single-cell and single-nucleus RNA-seq datasets, grouped by sequencing platform. These genes account for 23.8% of the 63 human-upregulated H_M4 genes identified in Extended Data Fig. 18. Violin plots show cellular expression distributions. Human, red; macaque, blue; mouse, green. Dataset sources are indicated on the x axes. Statistical annotations show within-platform comparisons with the reference human dataset using one-sided Wilcoxon rank-sum tests for higher human expression. *, *P* < 0.05; **, *P* < 0.01; ***, *P* < 0.001. *JUN*, *HES1* and *SOX9* are labelled in red. RG, radial glia.

**Extended Data Fig. 20.**
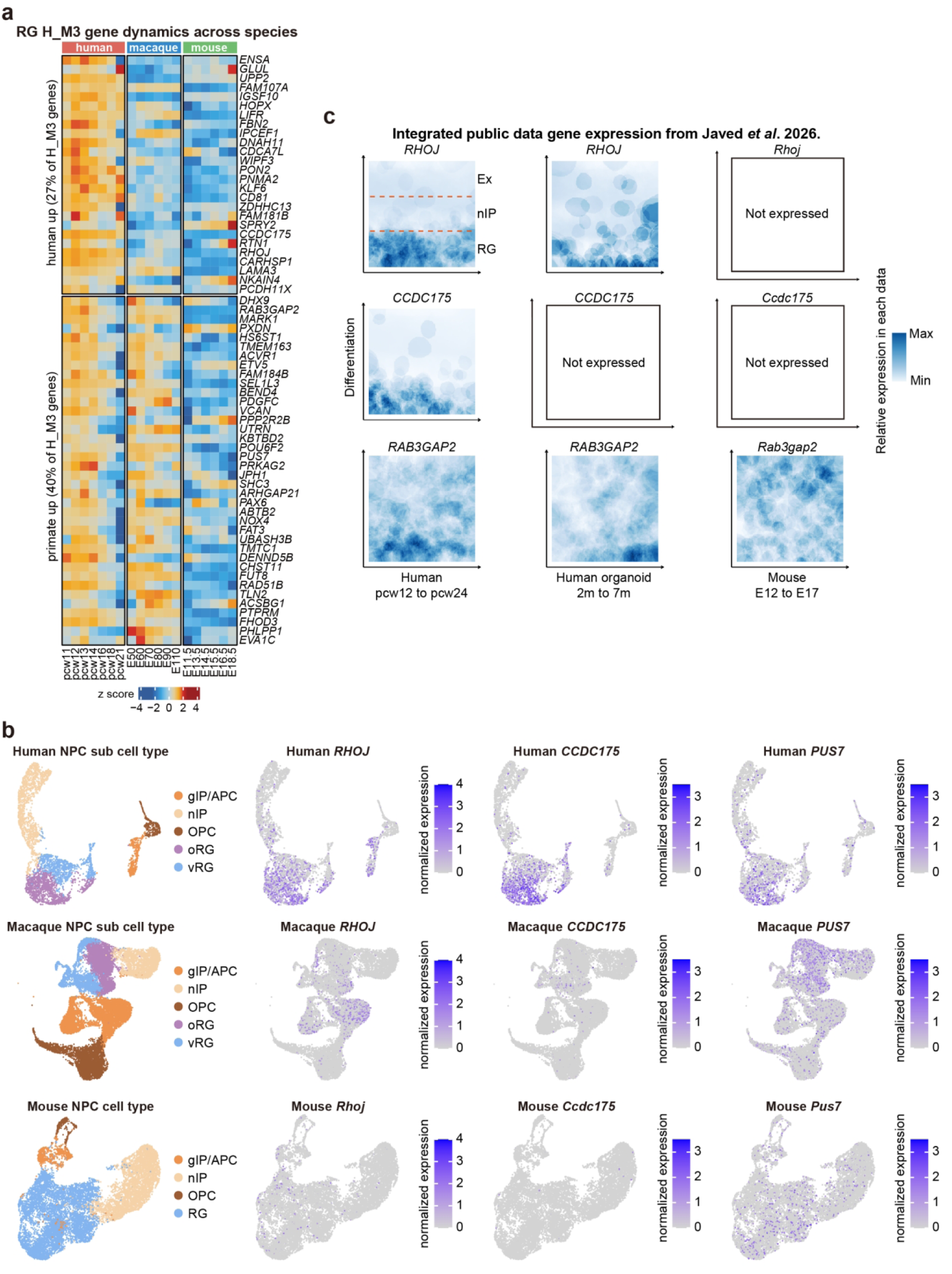
Cross-species expression profiles of H_M3 genes and candidate oRG markers. **a**, Temporal expression profiles of human-upregulated (27% of H_M3 genes) and primate-upregulated (40%) H_M3 genes in RG. Colours show gene-wise z scores across species and stages. **b**, Progenitor UMAPs for human, macaque and mouse, coloured by cell type (left) or normalized expression of *RHOJ*, *CCDC175* and *PUS7* (right). Mouse gene symbols denote the corresponding orthologs. **c**, Expression of *RHOJ*, *CCDC175* and *RAB3GAP2* in integrated public datasets from Javed *et al*. (2026): human fetal cortex (pcw12-24), human cortical organoids (2-7 months) and mouse cortex (E12-17). The vertical axis represents progression from RG towards neuronal states; colour intensity represents relative expression within each dataset. “Not expressed” denotes no detected expression in the displayed data. RG, radial glia; oRG, outer radial glia; vRG, ventricular radial glia; nIP, neuronal intermediate progenitors; gIP/APC, glial intermediate progenitors/astrocyte precursor cells; OPC, oligodendrocyte precursor cells; Ex, excitatory neurons; pcw, post-conception week; E, embryonic day.

**Extended Data Fig. 21.**
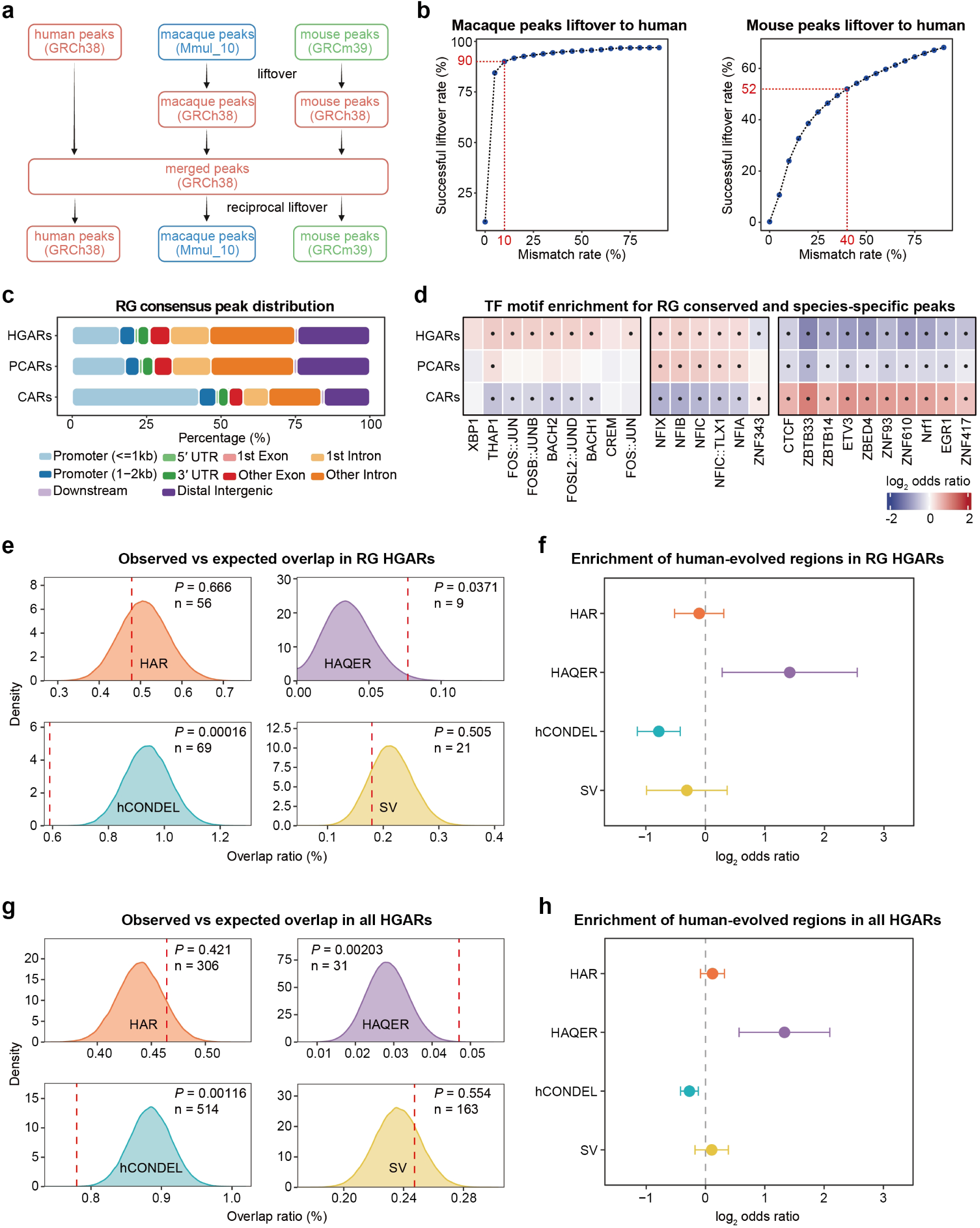
Cross-species accessible regions and their regulatory and evolutionary annotations. **a**, Reciprocal liftover workflow for defining accessible regions with one-to-one genomic correspondence among human GRCh38, macaque Mmul_10 and mouse GRCm39. **b**, Proportions of macaque and mouse peaks successfully mapped to human at different allowed mismatch rates. Red dashed lines indicate the selected tolerances of 10% for macaque and 40% for mouse. **c**, Genomic annotations of RG HGARs, PCARs and CARs. **d**, Enrichment of representative TF motifs, comparing each region class with the other two classes. Colours show log₂ Mantel–Haenszel odds ratios, stratified by GC content, region length and promoter overlap. Black dots indicate Benjamini–Hochberg-adjusted *P* < 0.05 from two-sided Cochran– Mantel–Haenszel tests, corrected across tested motifs and comparisons. **e**, Observed overlap of RG HGARs with human accelerated regions (HARs), human ancestor quickly evolved regions (HAQERs), regions associated with human-specific conserved deletions (hCONDELs) and human-specific structural variants (SVs), relative to conditional permutation distributions. Shaded distributions show overlap percentages from 100,000 permutations within strata defined by chromosome, region length and promoter overlap. Red dashed lines indicate observed overlap; n is the number of HGARs overlapping each annotation. Displayed *P* values are from two-sided permutation tests with Benjamini-Hochberg correction across the eight comparisons in **e** and **g**. **f**, Log₂ Mantel-Haenszel odds ratios for the comparisons in **e**, with 95% confidence intervals. The dashed line denotes no enrichment. **g**,**h**, Corresponding permutation distributions (**g**) and enrichment estimates (**h**) for the nonredundant union of HGARs across all ten analysed cell types. Backgrounds for **e**-**h** comprise the remaining cross-species-comparable accessible regions within the corresponding analysis scope. RG, radial glia; TF, transcription factor; HGAR, human-gained accessible region; PCAR, primate-conserved accessible region; CAR, conserved accessible region.

**Extended Data Fig. 22.**
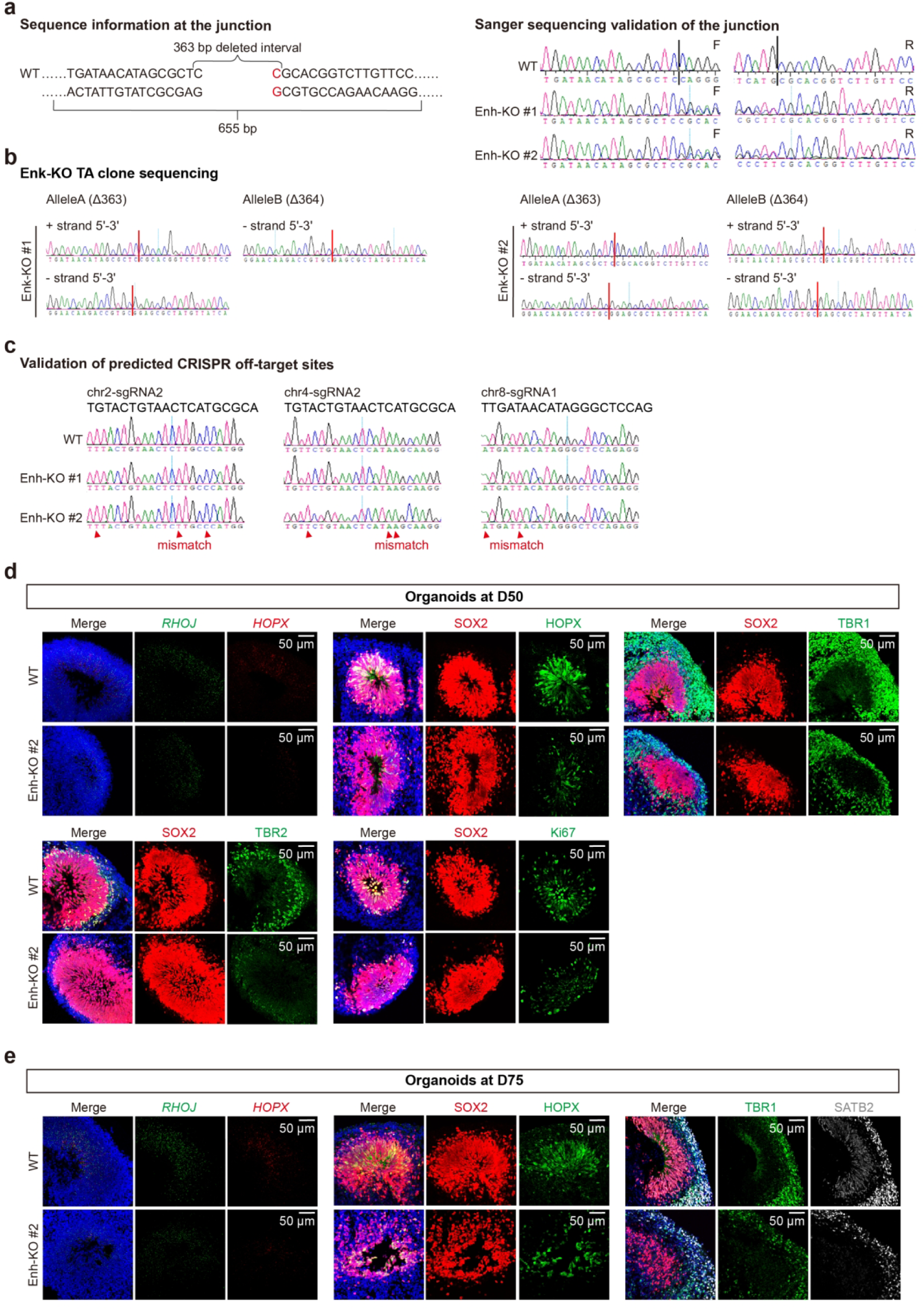
Validation of *RHOJ* enhancer-deletion clones and organoid phenotypes. **a**, Targeted enhancer sequence and Sanger sequencing of deletion junctions in WT, Enh-KO #1 and Enh-KO #2 cells. The targeted interval and PCR amplicon are indicated. F and R denote forward and reverse reads. **b**, Sanger sequencing of TA-cloned PCR products confirming biallelic deletion in both clones. Alleles A and B carry 363 bp and 364 bp deletions, respectively. Allele A/B support comprises 7/3 TA clones for Enh-KO #1 and 3/5 for Enh-KO #2. **c**, Sanger sequencing of predicted off-target sites on chromosomes 2, 4 and 8. No deletion-clone-specific sequence changes were detected at the examined sites. Arrowheads mark mismatches between the predicted target sequence and the sequenced locus. **d**, Representative D50 WT and Enh-KO #2 organoid images showing *RHOJ* and *HOPX* RNA-FISH, and immunofluorescence for SOX2 with HOPX, TBR1, TBR2 or Ki67. **e**, Representative D75 images showing RHOJ and HOPX RNA-FISH, SOX2/HOPX immunofluorescence and TBR1/SATB2 immunofluorescence. WT images in d and e are reproduced from the corresponding panels of Fig. 6 for comparison. Signal colours are indicated above each image; DAPI is blue. Scale bars, 50 µm (d,e). WT, wild type; D, day of organoid differentiation.

## Notes

### Competing Interest Statement

The authors have declared no competing interest.

